# Constitutive signal bias mediated by the human GHRHR splice variant 1

**DOI:** 10.1101/2021.08.20.457043

**Authors:** Zhaotong Cong, Fulai Zhou, Chao Zhang, Xinyu Zou, Huibing Zhang, Yuzhe Wang, Qingtong Zhou, Xiaoqing Cai, Qiaofeng Liu, Jie Li, Lijun Shao, Chunyou Mao, Xi Wang, Jihong Wu, Tian Xia, Lihua Zhao, Hualiang Jiang, Yan Zhang, H. Eric Xu, Xi Cheng, Dehua Yang, Ming-Wei Wang

## Abstract

Alternative splicing of G protein-coupled receptors has been observed, but their functions are largely unknown. Here, we report that a splice variant (SV1) of the human growth hormone releasing hormone receptor (GHRHR) is capable of transducing biased signal. Differing only at the receptor N terminus, GHRHR predominantly activates G_s_ while SV1 selectively couples to β-arrestins. Based on the cryo-electron microscopy structures of SV1 in the *apo* state or in complex with the G_s_ protein, molecular dynamics simulations reveal that the N termini of GHRHR and SV1 differentiate the downstream signaling pathways, G_s_ *vs*. β-arrestins. Suggested by mutagenesis and functional studies, it appears that GHRH-elicited signal bias towards β-arrestin recruitment is constitutively mediated by SV1. The level of SV1 expression in prostate cancer cells is also positively correlated with ERK1/2 phosphorylation but negatively correlated with cAMP response. Our findings imply that constitutive signal bias may be a mechanism that ensures cancer cell proliferation.

**Significance Statement:** The mechanism of functional changes induced by alternative splicing of GHRHR is largely unknown. Here, we demonstrate that GHRH-elicited signal bias towards β-arrestin recruitment is constitutively mediated by SV1. The cryo-electron microscopy structures of SV1 and molecular dynamics simulations reveal the different functionalities between GHRHR and SV1 at the near-atomic level, *i*.*e*., the N termini of GHRHR and SV1 differentiate the downstream signaling pathways, G_s_ *vs*. β-arrestins. Our findings provide valuable insights into functional diversity of class B1 GPCRs which may aid in the design of better therapeutic agents against certain cancers.

## Introduction

G protein-coupled receptors (GPCRs) are the largest superfamily of proteins in the body. They are almost expressed in every cell/tissue and transduce various signals to regulate a plethora of physiological functions (1). As a common and effective strategy to increase the functional diversity of the human genome, alternative splicing is often observed among GPCRs (2-5). The most common types of which include exon skipping, splice site selection and intron retention, resulting in deletion, exchange and insertion of receptor sequences, respectively (6). Although approximately 50% of GPCR genes are intronless, those that possess introns have the possibility to undergo alternative splicing, thereby generating subtype isoforms that may differ in ligand binding, receptor trafficking and signal transduction (7, 8). Some splice variants even display functional characteristics opposite to the canonical form (9).

GPCR splice variants often exhibit tissue-specific distribution and signaling characteristics that may impact disease pathology (10). For instance, Kahles *et al*. reported that alternative splicing events are more frequent in tumorous compared to normal tissues (11). Although splice variants of many GPCRs, such as growth hormone release hormone receptor (GHRHR) (12), thromboxane receptor (13), cholecystokinin-B receptor (14), secretin receptors (15) and somatostatin receptor (16), have been detected in various cancers, their biological significance is poorly understood. While alternative splicing of the C-X-C chemokine receptor 3 (CXCR3) was linked to β-arrestin recruitment (4), expression of GHRHR splice variants could be induced by hypoxic microenvironment in solid tumors leading enhanced glycolysis (17), suggesting that cancer-associated GPCR isoforms are not only a consequence of cellular adaptation, but also have an effect on malignancy.

We have previously revealed the structural basis of GHRHR activation and uncovered the detailed mechanism by which a naturally occurring mutation associated with isolated growth hormone deficiency leads to impaired GHRHR function (18). While the primary role of GHRHR is to stimulate growth hormone synthesis and secretion from the anterior pituitary somatotrophs upon GHRH binding (19), its ectopic expression in cancers has been studied extensively (20, 21). Both GHRH and GHRHR are present in lung (22), mammary (23), ovarian (24), endometrial (25), gastric (26), colorectal (27), ocular (28), prostatic (29, 30) and pancreatic (27) cancers. The cancer cell growth could be stimulated by exogenous GHRH and, conversely, inhibited by GHRHR antagonists (31). Among the four splice variants (SVs), SV1 possesses the greatest similarity to the full-length GHRHR and remains functional by eliciting cAMP signaling and mitogenic activity upon GHRH stimulation (32, 33). Due to its physical presence and bioactivity in cancer progression, SV1 is also called tumoral GHRHR that co-exists with pituitary GHRHR in most tumors. Compared with GHRHR, SV1 lacks a portion of the extracellular domain (ECD) because the first three exons are replaced by a fragment of intron 3, leading to the replacement of the first 89 amino acids of GHRHR with a distinct 25-amino acid sequence (12). Through functional diversity evaluation of almost all reported GPCR isoforms, Marti-Solano *et al*. found that the N-terminal splicing is the most frequently occurred structural variation and tends to alter ligand binding and/or signal transduction (34). In the case of SV1, changes of ligand binding affinities (35) and signaling properties are connected with its mitogenic effect (33, 36-40), but the underlying mechanism has yet to be elucidated.

Here, we report the cryo-electron microscopy (cryo-EM) structures of both the *apo* state and GHRH-bound SV1 in complex with G_s_ protein. Together with previously published GHRH– GHRHR–G_s_ complex structure (18), we are able to show the molecular details of ligand recognition and SV1 activation. In-depth investigations on SV1-mediated signal transduction unveiled a constitutively biased signaling pathway, thereby offering new insights into the role of alternative splicing of a class B1 GPCR in cancer cell proliferation.

## Results

### SV1 inhibits G_s_ activation

To better understand the functional outcome of alternative splicing of GHRHR, we evaluated the ability of SV1 to activate G_s_ upon stimulation by GHRH in HEK293T cells. HA signal peptide was fused to the Flag-tagged N terminus to rescue the cell surface expression level of SV1, which remained stable across all assays. G_s_ activation was assessed using split luciferase NanoBiT G-protein sensors to determine GHRH-induced decreases in luminescence on a time-course. For SV1, the G_s_ sensor gave a similar decrease in luminescence to GHRHR, suggesting that both caused the same G_s_ conformational change. However, GHRH concentration-response curves showed that the ability of SV1 to activate G_s_ was significantly impaired (Fig. 1*A*), as E_max_ and EC_50_ values were reduced (*SI Appendix* Table S1). G_s_-mediated cAMP accumulation was also drastically decreased (by almost 1000-fold) in cells expressing SV1 compared with that of GHRHR (Fig. 1*B* and *SI Appendix* Table S2). As expected, removal of the N-terminal 89 residues in GHRHR to imitate SV1 or deletion of the entire ECD led to diminished cAMP responses. In conjunction with the consequences of sequential deletion of 10 residues in the ECD of GHRHR, our results point to the importance of the N terminus in maintaining GHRHR function (Fig. 1*C*). GHRH-induced cAMP responses were subsequently measured in three prostate cancer cell lines expressing different levels of GHRHR and SV1. In line with previous findings (29, 30), LNCaP and 22Rv1 cells expressed high levels of GHRHR and SV1, respectively, while PC3 cells had low expression levels of both (Fig. 1*D* and *SI Appendix* Fig. S1*C*). It was found that LNCaP cells displayed the strongest cAMP response, whereas 22Rv1 and PC3 cells exhibited either markedly reduced or marginal cAMP responses (Fig. 1*E*).

**Figure 1.**
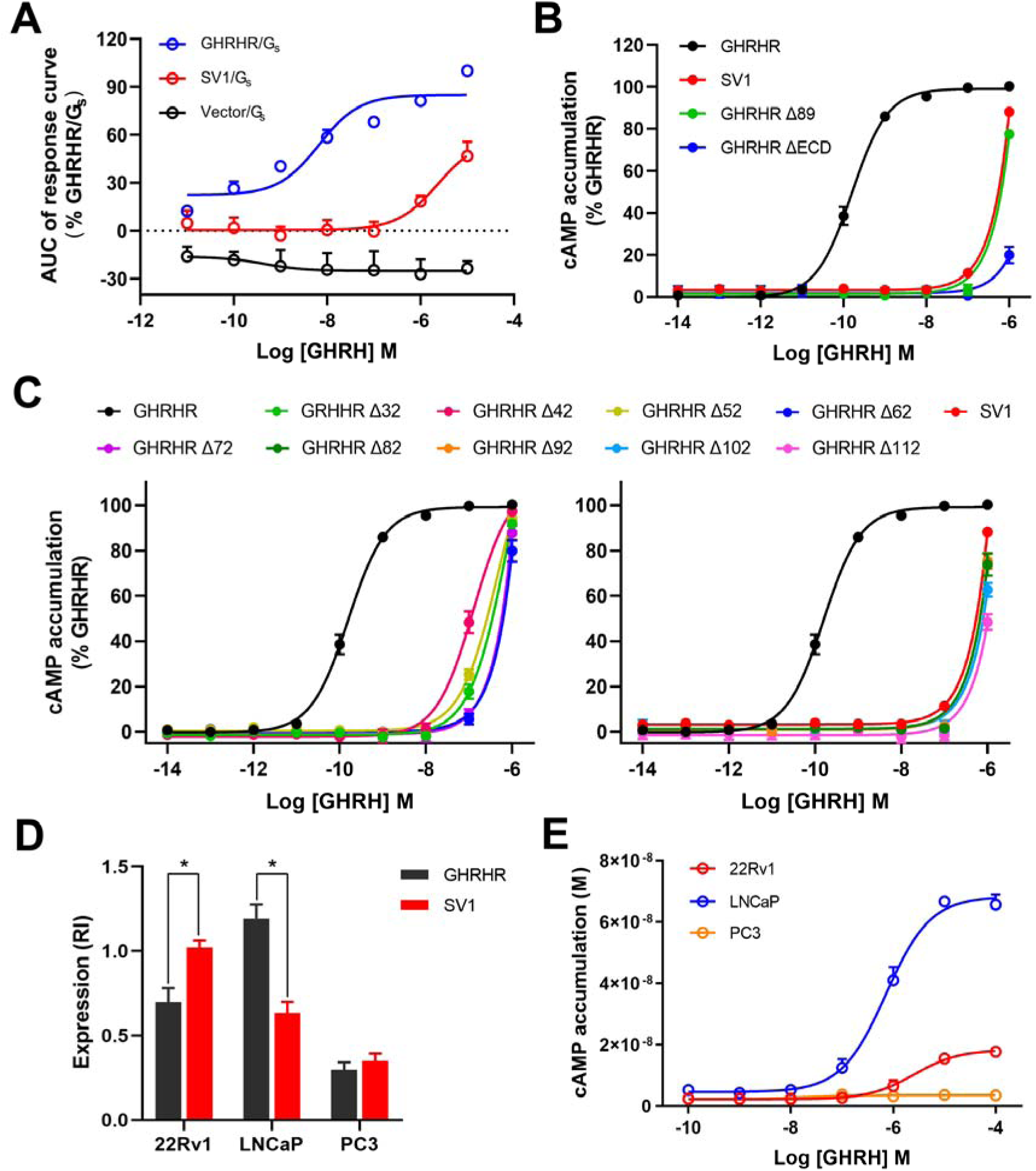
GHRH-induced G_s_ protein coupling and cAMP signaling mediated by GHRHR and SV1. (*A*) GHRH-induced conformational changes in the trimeric G_s_ protein. Concentration-response curves are expressed as AUC (area-under-the-curve) across the time-course response curve (0-25 min) for each concentration and normalized to GHRHR. (*B, C*) Concentration-response curves of cAMP accumulation at GHRHR and SV1. Comparison of SV1 with full-length or truncated GHRHR that lacks ECD or the first 89 residues (*B*). Comparison of SV1 with various N terminus truncated GHRHRs (*C*). (*D*) Expression of GHRHR and SV1 in prostate cancer cell lines. Protein levels were estimated as relative intensity (RI) compared to β-tubulin (loading control). (*E*) Concentration-response curves of cAMP accumulation in prostate cancer cells. Data shown are means ± S.E.M. of at least three independent experiments (*n* = 3-5) performed in quadruplicate. Δ, truncation; \**P* < 0.05; \*\**P* < 0.01.

### SV1 enhances β-arrestin recruitment

Since SV1 promotes cell proliferation and β-arrestin *per se* has pro-tumorigenic properties (41, 42), it is possible that SV1 may potentially behave as a β-arrestin-biased variant that facilitates cancer cell growth. To test this hypothesis, we measured GHRH-induced β-arrestin recruitment and observed that SV1 indeed increased both β-arrestin 1 and β-arrestin 2 recruitment by about 18% and 30%, respectively, compared with that of GHRHR (Fig. 2*A*). β-arrestin 1/2 recruitment was also enhanced following truncation of the first 89 residues or the entire ECD -; it was not affected if the deletion was made before the first 82 residues (Fig. 2*B*), suggesting that this biased signaling is caused by a structural change in the ECD of GHRHR. We next measured ERK1/2 phosphorylation (pERK1/2) upon GHRH stimulation in HEK293T cells expressing GHRHR or SV1. In cells expressing SV1, GHRH induced a stronger pERK1/2 response which was about 40% higher than that of GHRHR at the peak response time (Fig. 2*C* and *SI Appendix* Fig. S1*A*). To examine if the ERK phosphorylation was GHRHR- or SV1-dependent, the cells were also treated with a GHRHR antagonist, MIA-602, before GHRH stimulation. Inhibition of pERK1/2 by MIA-602 indicates that GHRH could specifically activate this signaling pathway (Fig. 2*C* and *SI Appendix* Fig. S1*A*). Consistent with previous findings showing that SV1 promotes cell proliferation via pERK1/2 pathway (30, 40), we found that SV1 expression is correlated with the cell cycle. One μM GHRH augmented the number of cells in G_2_/M phase (increased from 22.8% to 37.7%) but diminished that corresponding to G_1_ phase (decreased from 46.6% to 32.8%) (*SI Appendix* Fig. S1*C*). Among the three prostate cancer cell lines, only 22Rv1 that expresses a high level of SV1 displayed a markedly stronger and longer pERK1/2 response which could be inhibited by MIA-602 (Fig. 2*D* and *SI Appendix* Fig. S1*B*).

**Figure 2.**
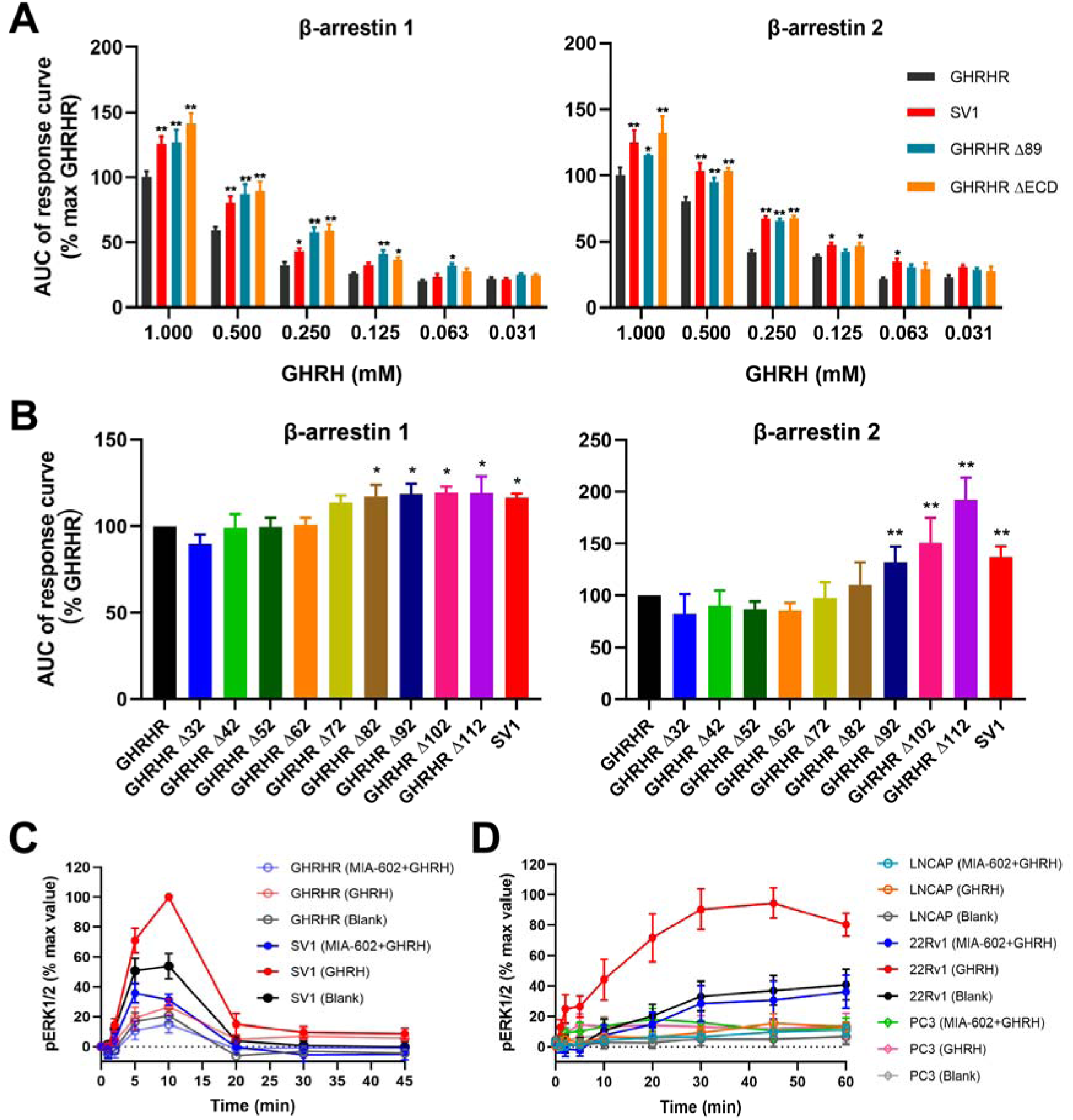
GHRH-induced β-arrestin recruitment and pERK1/2 signaling mediated by GHRHR and SV1. (*A, B*) β-arrestin recruitment by GHRHR and SV1. Comparison of SV1 with full-length or truncated GHRHR that lacks the entire ECD or the first 89 residues (*A*). Comparison of SV1 and various N terminus truncated GHRHRs. The assay was initiated by 250 μM GHRH (*B*). The response was calculated as AUC (area-under-the-curve) across the full kinetic trace. *P < 0.01 and **P < 0.001 compared with GHRHR; Δ, truncation. (*C, D*) Time-course of ERK1/2 activation. The assay was initiated by 1 μM GHRH and inhibition was achieved by 4 μM MIA-602 in HEK293T cells expressing GHRHR or SV1 (*C*) and prostate cancer cell lines (*D*). Data shown are means ± S.E.M. of at least four independent experiments (*n* = 4-6) performed in duplicate. \**P* < 0.05; \*\**P* < 0.01.

### Structure comparison between GHRHR and SV1

To obtain a stable SV1 complex for cryo-EM study, we removed 18 amino acids at the C terminus and employed the NanoBiT tethering strategy (18, 43) to further stabilize the assembly of SV1 with G_s_ heterotrimer (*SI Appendix* Fig. S2). The structures of GHRH–SV1–G_s_ and SV1–G_s_ (*apo*) complexes were determined by single-particle cryo-EM at global resolutions of 3.3Å and 2.6Å, respectively (Fig. 3, *SI Appendix* Fig. S3-4 and *SI Appendix* Table S3). The high-quality density maps allowed unambiguous building for receptor residues L55^1.29b^ to H328^8.59b^ (class B GPCR numbering in superscript (44)), GHRH (residues Y1^P^-A19^P^), and most residues of Nb35 and Gαβγ subunits except the α-helical domain (AHD) of Gα_s_. The majority of amino acid side chains, except for residues C131^ECL1^-S132^ECL1^ as well as P249^ICL3^-H255^6.30b^ in the *apo* SV1–G_s_ complex and residues P249^ICL3^-Q255^6.32b^ in the GHRH–SV1–G_s_ complex, were well resolved in the final models (*SI Appendix* Fig. S5).

**Figure 3.**
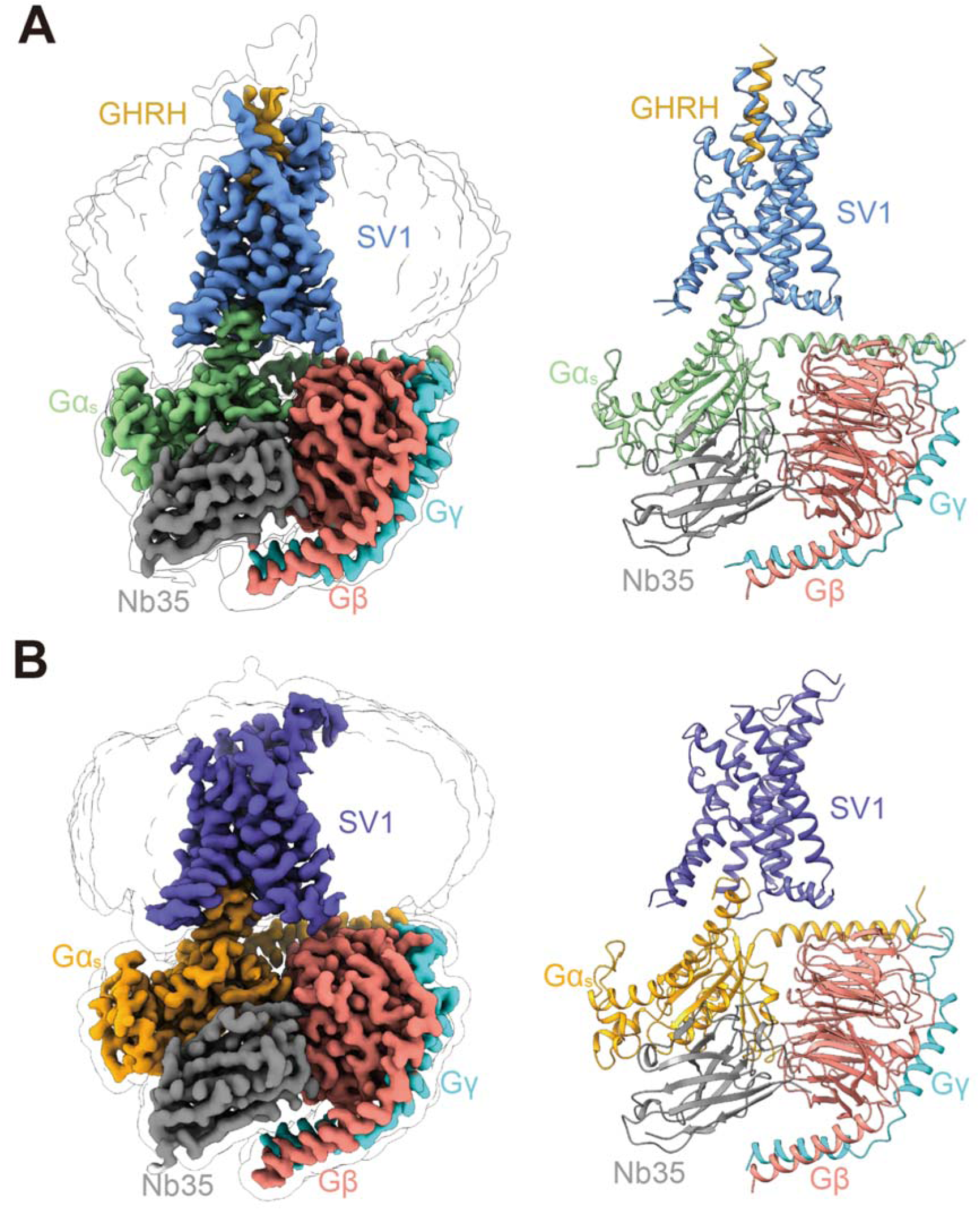
Overall structures of GHRH–SV1–Gα_s_ and *apo* SV1–Gα_s_ complexes. (*A*) Orthogonal views of the density map (left) and the model (right) for the GHRH–SV1–Gα_s_–Nb35 complex. SV1, GHRH, Gα_s_, Gβ, Gγ and Nb35 are colored cornflower blue, gold, light green, salmon, cyan and gray, respectively. (*B*) Orthogonal views of the density map (left) and the model (right) for the *apo* SV1–Gα_s_–Nb35 complex. SV1, Gα_s_, Gβ, Gγ and Nb35 are colored slate blue, orange, salmon, cyan and gray, respectively. The structures are shown in cartoon representation.

In the GHRH–SV1–G_s_ complex structure, the bundle of seven transmembrane (TM) helices adopts highly similar conformations to that of the GHRH–GHRHR–G_s_ complex (18) with a Cα root-mean-square deviation of 0.5Å (Fig. 4*A*). This was expected, considering the amino acid sequences of transmembrane domain (TMD) are identical between SV1 and GHRHR. However, the interactions between ECD and GHRH are remarkably different in the two complexes. In the GHRH–SV1–G_s_ complex, its ECD does not stabilize the binding by GHRH (Fig. 4*A*), while that of GHRHR has rich interactions with GHRH involving residues L34^ECD^, L62^ECD^, F82^ECD^ and F85^ECD^ (Fig. 4*B*). Substituting any or all of these four ECD residues with alanine reduced GHRHR elicited cAMP responses (*SI Appendix* Fig. S6*B* and Table S2) but enhanced β-arrestin 1/2 recruitments (*SI Appendix* Fig. S6*A*), similar to SV1. These findings underscored the importance of the N-terminal ECD residues in determining signal bias upon GHRH stimulation. Notably, without stable interactions with residues in the SV1 ECD, the C terminus of GHRH is highly flexible. Consequently, the resolvable region of GHRH in the GHRH–SV1–G_s_ complex is ten-residue shorter at the C terminus than that in the GHRH–GHRHR–G_s_ complex (Fig. 4*A*). In the GHRHR complex structure, GHRH has a slightly larger tilting angle than that in the SV1 complex structure due to its interactions with the ECD of GHRHR (*SI Appendix* Fig. S7). Nevertheless, due to resembling TMD conformations, the N terminus of GHRH binds to the orthosteric pocket of SV1 with an orientation similar to that in the GHRH–GHRHR–G_s_ complex (Fig. 4*A*).

**Figure 4.**
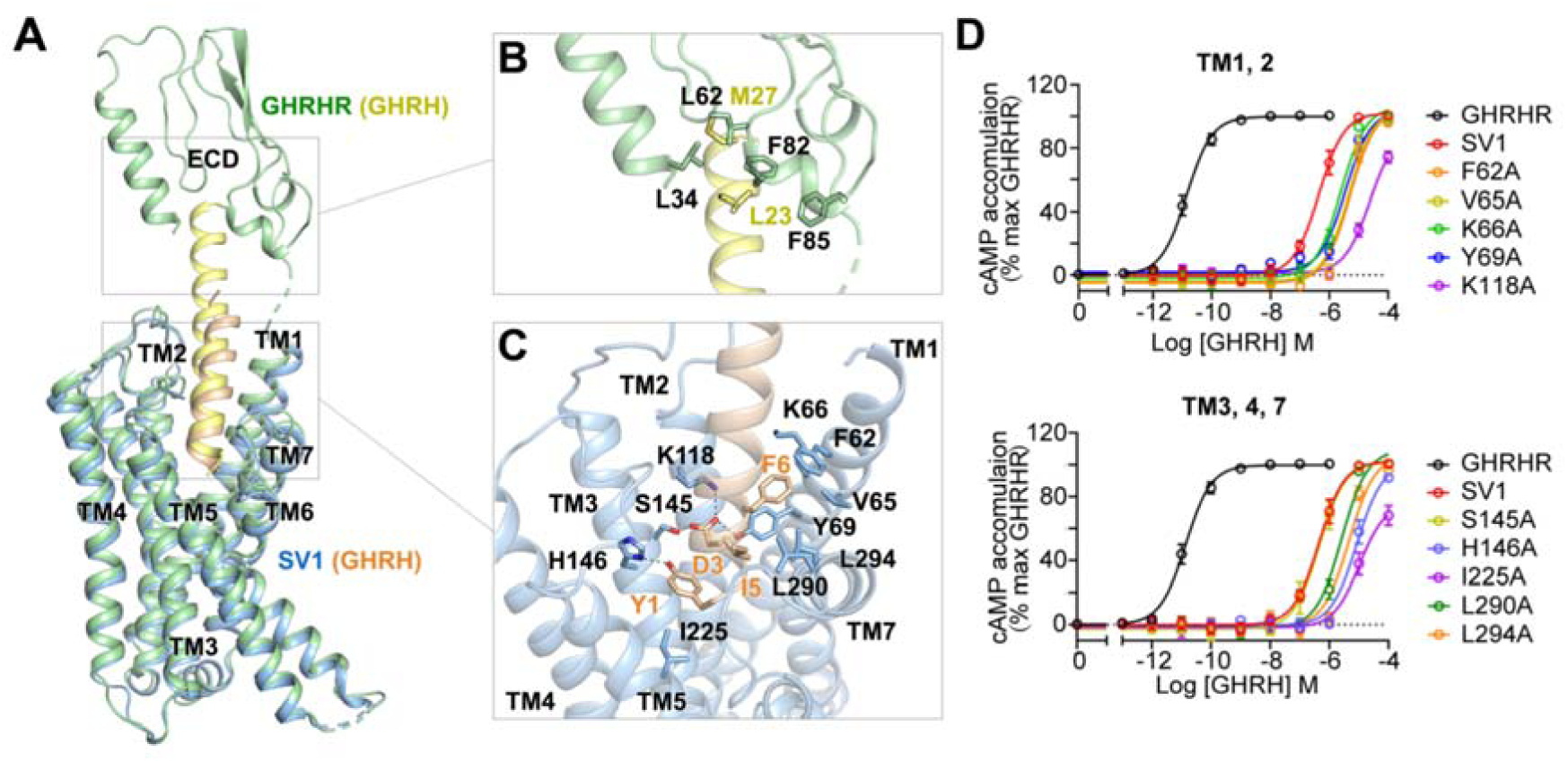
Structural comparison between SV1 and GHRHR. (*A*) Comparison between the cryo-EM structures of GHRH–SV1–G_s_ and GHRH–GHRHR–G_s_ complexes. Receptors and GHRH are shown in cartoon: GHRHR is colored in green, SV1 in blue, GHRH in wheat and yellow. G_s_ is omitted for clarity. (*B*) Detailed interaction between GHRH (yellow) and the ECD of GHRHR (green). Key residues are shown as sticks. (*C*) Detailed interaction between GHRH (wheat) and the peptide-binding pocket of SV1 (blue). Salt bridges and hydrogen bonds are shown as dash lines. (*D*) Effects of mutations in the peptide-binding pocket of SV1 on cAMP accumulation. Data shown are means ± S.E.M. of four independent experiments (*n* = 4) conducted in quadruplicate.

The GHRH–SV1–G_s_ complex structure shows that GHRH binds to SV1 through a continuous interacting network involving TMD helices (TMs 1-4, and TM7) (Fig. 4*A* and *C*). The N terminus of GHRH deeply inserts into the receptor core. Y1^P^ of GHRH forms hydrogen bonds with H146^3.37b^ and hydrophobic interactions with I225^5.43b^. D3^P^ makes salt bridges with K118^2.67b^, which is further strengthened by hydrogen bonding with Y69^1.43b^ and S145^3.36b^. I5^P^ has van der Waals interactions with two TM7 residues, *i*.*e*., L290^7.35b^ and L294^7.39b^. F6^P^ builds extensive hydrophobic contacts with F62^1.36b^, V65^1.39b^ and K66^1.40b^. Impairing these contacts dramatically decreased the potency of GHRH-induced cAMP accumulation mediated by SV1 (Fig. 4*D* and *SI Appendix* Table S4), suggesting essential roles of these residues in ligand recognition and receptor activation. Meanwhile, binding of GHRH yields to an extended helical conformation of TM1 and inward movements of ECLs 1-2 to the peptide-binding pocket of SV1 (*SI Appendix* Fig. S8).

### Interaction between SV1 and β-arrestin 1

To gain insights into the molecular mechanism by which β-arrestins bind to GHRHR and SV1, we performed molecular dynamics (MD) simulations of arrestin-bound receptors. Because previous structural studies have reported a few β-arrestin 1-bound GPCR complexes but there is none complexed with β-arrestin 2, we constructed two simulation systems for GHRHR and SV1 bound to β-arrestin 1, respectively. In the GHRHR simulations, the ECD constantly bound to GHRH (Fig. 5*A, B* and *SI Appendix* Fig. S9*A*). Multiple hydrophobic ECD residues (F30^ECD^, I31^ECD^, L34^ECD^, L62^ECD^, F81^ECD^, F82^ECD^ and F85^ECD^) of GHRHR frequently interacted with GHRH to stabilize its binding during simulations (Fig. 5*C* and *SI Appendix* Fig. S9*A*). On the contrary, in the SV1 system, the short ECD of the receptor did not stably interact with the peptide (Fig. 5*A* and *B*), resulting in an outward movement of GHRH at the extracellular side (Fig. 5*B*). The average minimal distance between GHRH and the bottom of the peptide-binding pocket of SV1 was 6.5 ± 1.6Å, approximately 3Å longer than that of the GHRHR system (3.4 ± 0.2Å). In the GHRHR simulations, GHRH stably bound to the peptide-binding pocket and frequently interacted with ECL2 (Fig. 5*D*). Notably, the peptide residue R11^P^ could form a salt bridge with D274^ECL2^ in the GHRHR system. These observations suggest that the ECD of GHRHR binds to GHRH to stabilize its orientation in the peptide-binding pocket and further enhances its interactions with ECL2 at the extracellular interface. However, the ECD of SV1 is too short to stabilize the orientation of GHRH that failed to interact with ECL2 (Fig. 5*D*).

**Figure 5.**
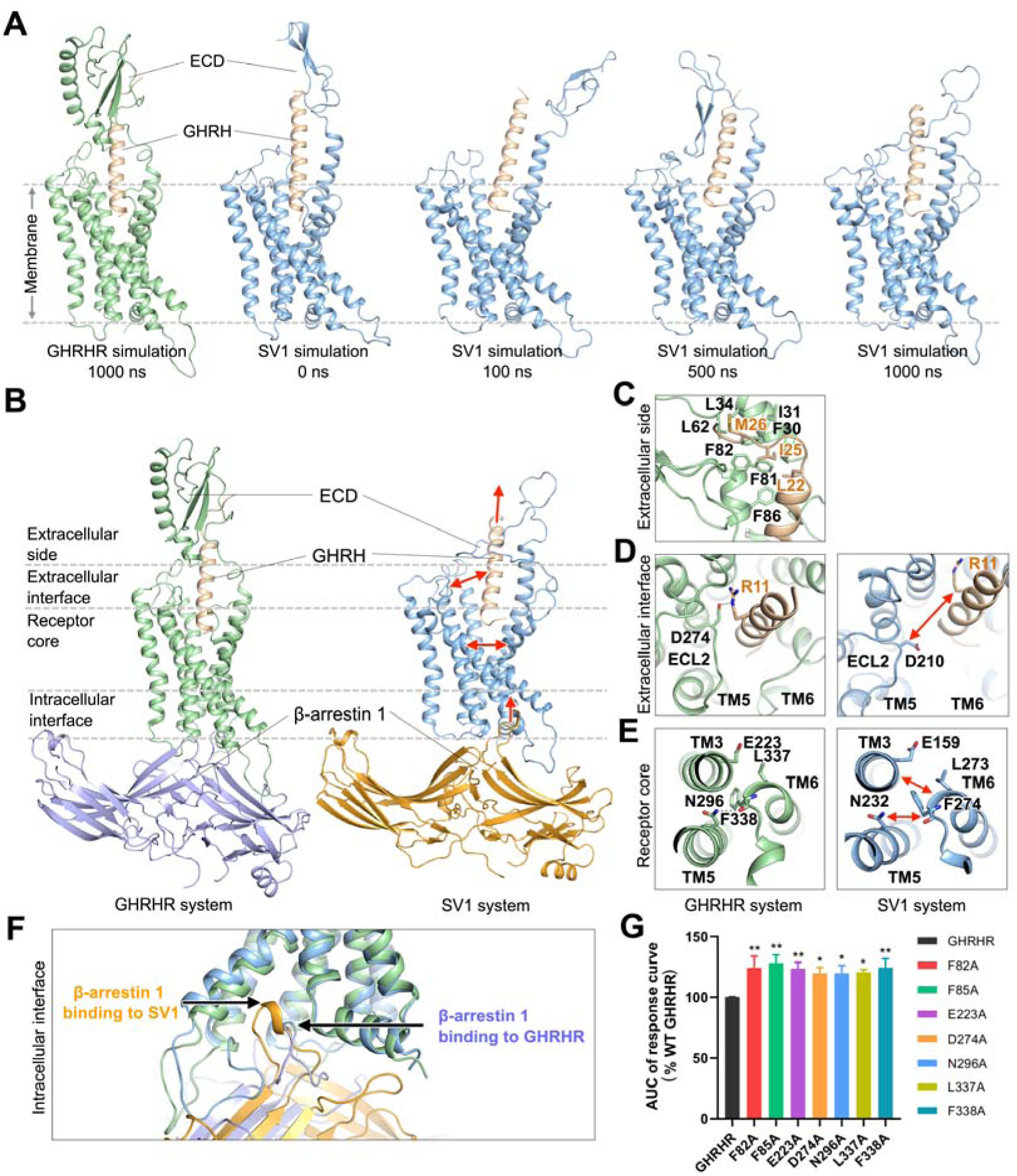
β-arrestin 1 binding to GHRHR and SV1. (*A*) Distinct ECD conformations of GHRHR and SV1 during simulations. Receptors and GHRH are shown in cartoon: GHRHR is colored in green, SV1 in blue and GHRH in wheat. β-arrestin 1 is omitted for clarity. (*B*) Representative simulation snapshots from GHRHR (left) and SV1 (right) systems. Gray dash lines split the different regions of a receptor. (*C*) A representative simulation snapshot showing key interactions between GHRH (wheat) and the ECD of GHRHR (green). Key residues are shown as sticks. (*D*) Representative simulation snapshots showing the extracellular interfaces of GHRHR (left) and SV1 (right). A salt bridge of GHRHR (green) is shown as a black dash line. (*E*) Representative simulation snapshots showing the receptor cores of GHRHR (left) and SV1 (right). A hydrogen bond of GHRHR (green) is shown as a black dash line. (*F*) Binding of β-arrestin 1 to GHRHR (green) and SV1 (blue) at the intracellular side in simulations. (*G*) β-arrestin 1 recruitment by GHRHR and its mutants. The assay was stimulated by 250 μM GHRH. Data shown are means ± S.E.M. of at least three independent experiments (*n* = 3-6) performed in duplicate; \**P* < 0.05, \*\**P* < 0.01.

As a loop linking TM5 helix, ECL2 contributes to determining its orientation. Interacting with GHRH, the ECL2 of GHRHR stayed adjacent to the peptide-binding pocket and pulled TM5 helix toward the pocket center, which contributed to a compact buddle of helices TM3, TM5 and TM6 (Fig. 5*D*). In particular, TM5 residue N296^5.50b^ formed a hydrogen bond with the backbone oxygen of TM6 residue F338^6.49b^, and TM6 residue L337^6.48b^ stably interacted with the hydrophobic part of TM3 residue E223^3.50b^ at the receptor core of GHRHR (Fig. 5*E*). In the SV1 simulations, ECL2 did not stably interact with GHRH and wandered around the extracellular interface, thereby disrupting the interactions among helices TM3, TM5 and TM6 (Fig. 5*E*). The average minimal backbone distance between TM3 and TM6 increased to 7.3 ± 0.7Å, and the backbone distance between TM5 and TM6 increased to 7.1 ± 0.5Å. No hydrogen bonding was formed between TM5 and TM6 at the receptor core during the SV1 simulations (Fig. 5*E*). The hydrophobic interaction between the TM3 glutamic acid (E159^3.50b^) and the TM6 leucine (L273^6.48b^) of SV1 was also missing (Fig. 5*E*). As the inter-helical distance among TM3, TM5 and TM6 increased, the volume of arrestin-binding pocket of SV1 enlarged to 2441.3 ± 99.1Å^3^ at the intracellular side, approximately 300Å^3^ larger than that of GHRHR (*SI Appendix* Fig. S9*B*). Consequentially, β-arrestin 1 had a deeper insertion towards the receptor core of SV1 (Fig. 5*F*). Compared with the GHRHR system, the insertion of β-arrestin 1 into SV1 was approximately 3Å deeper along the membrane (z-axis in a simulation system). During the SV1 simulations, the finger loop of β-arrestin 1 exhibited more interactions with the receptor. Particularly, L68 of β-arrestin 1 stably interacted with the TM3 residues L163^3.54b^ and L166^3.57b^ of SV1, but did not interact with TM3 of GHRHR (*SI Appendix* Fig. S9*C*). These findings indicate that SV1 might provide a more favorable binding interface for β-arrestin 1, compared with GHRHR. Collectively, with a short ECD, SV1 might be unable to stabilize the orientation of GHRH and therefore could not maintain the interaction between GHRH and ECL2, leading to a loop buddle of helices TM3, TM5 and TM6 and an enlarged favorable arrestin-binding pocket. For GHRHR, the full-length ECD stabilizes GHRH and facilitates GHRH-binding to ECL2, which might make a compact buddle of helices TM3, TM5 and TM6 and a less favorable pocket for arrestin binding. To validate this hypothesis, we designed single-point mutations in the crucial domain of GHRHR mentioned above. Substituting a residue in these domains (F82^ECD^ and F85^ECD^ in ECD, D274^ECL2^ in ECL2, E223^3.50b^ in TM3, N296^5.50b^ in TM5, and L337^6.48b^ and F338^6.49b^ in TM6) with alanine significantly increased the capability of GHRHR to recruit β-arrestin 1 (Fig. 5*G* and *SI Appendix* Fig. S6*C*), fully supporting our hypothesis.

## Discussion

Although both GHRHR and SV1 are present in prostatic (45), breast (46), gastric (27), ovarian (47), pancreatic (37), lung (48), esophageal (17), oral (38, 49) and skin cancers (50), SV1 possess stronger mitogenic activities (36, 39, 40). A recent study demonstrates that alternative splicing of GHRHR, promoted by hypoxic microenvironment in solid tumors, is actually a cellular adaptation mechanism that induces cancer cell proliferation and migration (17). As a splice variant of GHRHR, SV1 only differs by 89 amino acids at its N terminus but causes a distinct biased signaling which may involve a complex regulatory mechanism associated with cancer development.

Integrating structure determination with functional studies, we reveal that splicing encoded structural difference in the ECD of GHRHR can alter its signaling preference. Compared with GHRHR, which predominantly couples to G_s_, SV1 preferentially activates β-arrestins. Removal of the N-terminal 89 residues in GHRHR to imitate the splicing or deletion of the entire ECD led to diminished cAMP responses and enhanced β-arrestin 1/2 recruitments, pointing to the importance of the N terminus in the constitutive signal bias. To uncover the molecular basis of this phenomenon, we solved the cryo-EM structures of SV1 in the *apo* state and in complex with G_s_ protein. In addition, we performed MD simulations to study conformational changes of SV1 mediated by β-arrestin 1 that may subsequently differentiate the downstream signaling pathways. The positive correlation of pERK1/2 and the negative correlation of cAMP accumulations with SV1 expression levels in prostate cancer cells implying a linkage with cancer. Therefore, modification of GHRHR by splicing at the N terminus results in biased arrestin signaling that might be advantageous to tumor cells.

Based on the SV1 structural information, MD simulations and mutagenesis analysis, we found that alternation of the ECD might change the intracellular interface that binds to β-arrestins. In the case of GHRHR, its full-length ECD stably constrains the orientation of GHRH and facilitates the interaction between ECL2 and GHRH, which might lead to a compact buddle of TMD and a less favorable intracellular interface for β-arrestins. In contrast, with a shorter and flexible ECD, SV1 does not stabilize the interaction between GHRH and ECL2, but possibly contributes to a loop buddle of helices TM3, TM5 and TM6 and a binding pocket permitting optimal insertion of the finger loop of β-arrestins. The ECD residues efficiently regulate signal bias via interacting with ligand. Particularly, four hydrophobic residues of GHRHR (L34^ECD^, L62^ECD^, F82^ECD^ and F85^ECD^) bound to GHRH are identified as key ECD elements that determine the nature of downstream signals. Substituting them with alanine significantly decreased cAMP responses (*SI Appendix* Fig. S6*B*). F82A and F85A mutants of GHRHR enhanced β-arrestin 1/2 recruitment, similar to that seen with SV1 (*SI Appendix* Fig. S6*A*) and consistent with the outcome of ECD truncation (Figs. 1, 2).

Alternation of N termini has been reported to modulate ligand binding and/or activity of a number of GPCRs (34). For instance, the N-terminal residues of CXCR3 determine its selectivity to a particular effector, *i*.*e*., β-arrestin 2 (4). In addition to ECD, ECL2 may also play an essential role in transducing signal from the extracellular side to the receptor core. Substitution of D274^ECL2^ with an alanine enhances β-arrestin signaling (Fig. 5*G*). The interaction between ECL2 and parathyroid hormone (PTH) facilitates β-arrestin recruitments by PTH receptor type 1 (51). In the receptor core, TMD helices rearrange in favor of interacting with G protein or β-arrestins. Disruption of which by substituting E223^3.50b^, N296^5.50b^, L337^6.48b^ and F338^6.49b^ in of GHRHR with alanine caused a biased β-arrestin signaling (Fig. 5*G*). The proposed molecular mechanism unveils the potential roles of ECD and ECL2, TMD in signal bias, which might extend to other class B GPCRs.

The signal bias of GPCRs can be classified as ligand bias and receptor bias (52). As a principal component to initiate signaling, a receptor itself is capable of constitutively biasing downstream signal transduction through genetic variations, including splice variant. A number of naturally occurring mutations were found to alter signaling pathways of GPCRs (52). For example, substitution of a TM6 residue of α1-adrenergic receptor led to constitutive G protein activity (53); a leucine-to-glutamine mutation in the TM3 helix of cysteinyl-leukotriene receptor 2 strongly drove G_q/11_ signaling (54); and mutations in the C terminus of several class A GPCRs, including apelin receptor and neuropeptide Y4 receptor, diminished β-arrestin recruitments (55, 56). Although many genetic variations of GPCRs have been detected (34), the mechanisms governing signal bias are poorly understood. In this work, we show that a splice variant strongly drives β-arrestin recruitment by averting the canonical signaling which is biased to G_s_ pathway. This structural alternation not only allows normal cells to function via a full-length receptor but also permits cancer cell to proliferate through a splice variant upon stimulation by the same endogenous ligand. Our findings thus provide new insights into functional diversity of class B1 GPCRs and offer valuable information to design of better therapeutics against certain cancer.

## Materials and Methods

### Cell culture

*Spodoptera frugiperda* (*Sf*9) insect cells (Expression Systems) were grown in ESF 921 serum-free medium (Expression Systems) at 27°C and 120 rpm. HEK293T cells were purchased from American Type Culture Collection (ATCC), cultured in Dulbecco’s modified Eagle’s medium (DMEM, Life Technologies) supplemented with 10% fetal bovine serum (FBS, Gibco) and maintained in a humidified chamber with 5% CO_2_ at 37°C. The PC3 prostate cancer cell line (ATCC) was cultured in F12 medium supplemented with 10% FBS and maintained in a humidified chamber with 5% CO_2_ at 37°C. 22Rv1 and LNCaP human prostate cancer cells (ATCC) were cultured in RPMI medium 1640 supplemented with 10% FBS and maintained in a humidified atmosphere containing 5% CO_2_ at 37°C.

### Constructs of SV1 and G_s_ heterotrimer

To facilitate the expression and purification, human wild-type (WT) SV1 gene with the hemagglutinin (HA) signal peptide at its N terminus, eighteen amino acids (A342-C359) truncation and a TEV protease cleavage site followed by a double maltose-binding protein-(MBP) tag at its C terminus was cloned into the pFastBac vector (Invitrogen). To obtain a SV1–G_s_ complex with good homogeneity and stability, we used the NanoBiT tethering strategy (18, 43, 57), in which the C terminus of rat Gβ1 was linked to HiBiT subunit and the C terminus of SV1 was directly attached to LgBiT subunit with a 15-amino acid polypeptide (GSSGGGGSGGGGSSG) linker. A dominant-negative human Gα_s_ (DNGα_s_) was generated by site-directed mutagenesis as previously described (58) to limit G protein dissociation. An engineered G_s_ construct (G112) was designed based on mini-G_s_ (59, 60) that was employed in the determination of A_2A_R-mini-G_s_ crystal structure (61). It was used to purify the *apo* state SV1–G_s_ complex. By replacing N-terminal histidine tag (His6) and TEV protease cleavage site with the N-terminal eighteen amino acids (M1-M18) of human G_i1_, the chimeric G_s_ was capable of binding to scFv16, which was used to stabilize the GPCR-G_i_ or -G_11_ complexes (62, 63). Additionally, replacement of the GGSGGSGG linker at position of original Gα_s_ α-helical domain (AHD, V65-L203) with that of human G_i1_ (G60-K180) provided the binding site for Fab_G50, an antibody fragment which was used to stabilize the rhodopsin-G_i_ complex (64). Furthermore, three mutations (G226A, L272D and A366S) were also incorporated by site-directed mutagenesis as previously described to further increase the dominant-negative effect by stabilizing the Gαβγ heterotrimer (59). These modifications enabled the application of different nanobodies or antibody fragments to stabilize the receptor-G_s_ complex, although Nb35 was solely used during SV1-G_s_ complex formation and stabilization in this study. The engineered G_s_ has also been employed and validated in the cryo-EM structure determination of the vasopressin V2 receptor–G protein complex (59).

### Expression and purification of nanobody 35

The nanobody 35 (Nb35) with a C-terminal histidine tag (His6) was expressed in *E. coli* BL21 (DE3) bacteria and cultured in TB medium supplemented with 2 mM MgCl_2_, 0.1% (w/v) glucose and 50 μg/mL ampicillin to an OD_600_ value of 1.0 at 37°C. The cultures were then induced by 1 mM IPTG and grown for 5 h at 37°C. Cells were harvested by centrifugation (4,000 rpm, 20 min) and Nb35 protein was extracted and purified by nickel affinity chromatography as previously described (65). Eluted protein was concentrated and subjected to a HiLoad 16/600 Superdex 75 column (GE Healthcare) pre-equilibrated with buffer containing 20 mM HEPES, pH 7.5 and 100 mM NaCl. The monomeric fractions supplemented with 30% (v/v) glycerol were flash frozen in liquid nitrogen and stored in -80°C until use.

### Expression and purification of the SV1-G_s_ complex

*Sf*9 insect cells were cultured at a density of 3 × 10^6^ cells per mL and co-infected with SV1-15AA-LgBiT, DNGα_s_ or engineered G_s_, Gβ1-15AA-peptide 86 and Gγ2 baculoviruses at a 1:1:1:1 ratio. The cells were then harvested by centrifugation 48 h post-infection and stored in -80°C for future use. The frozen cells were thawed on ice and resuspended in lysis buffer containing 20 mM HEPES, pH 7.5, 100 mM NaCl, 10% (v/v) glycerol, 10 mM MgCl_2_, 5 mM CaCl_2_, 1 mM MnCl_2_, 100 μM TCEP (Sigma-Aldrich) and supplemented with EDTA-free protease inhibitor cocktail (Bimake). Cells were lysed by dounce homogenization and complex formation was initiated in the presence of 10 μg/mL Nb35, 25 mU/mL apyrase (Sigma-Aldrich) and 20 μM GHRH (GL Biochem) for 1.5 h at room temperature (RT). The membrane was then solubilized by adding 0.5% (w/v) lauryl maltose neopentyl glycol (LMNG, Anatrace) and 0.1% (w/v) cholesterol hemisuccinate (CHS, Anatrace) for 2 h at 4°C. After centrifugation at 30,000 rpm for 30 min, the sample was clarified and the supernatant was incubated with amylose resin (NEB) for 3 h at 4°C. After incubation, the resin was collected by centrifugation (600 *g*, 10 min) and loaded to a gravity flow column, followed by five column volumes wash of buffer A containing 20 mM HEHES, pH 7.5, 100 mM NaCl, 10% (v/v) glycerol, 5 mM MgCl_2_, 1 mM MnCl_2_, 2 μM GHRH, 25 μM TCEP, 0.1% (w/v) LMNG and 0.02% (w/v) CHS and fifteen column volumes wash of buffer B containing 20 mM HEHES, pH 7.5, 100 mM NaCl, 10% (v/v) glycerol, 5 mM MgCl_2_, 1 mM MnCl_2_, 2 μM GHRH, 25 μM TCEP, 0.03% (w/v) LMNG, 0.01% (w/v) glyco-diosgenin (GDN, Anatrace) and 0.008% (w/v) CHS. The bound samples were incubated with His-tagged TEV protease (customer-made) overnight at 4°C in buffer B. The flow through was collected next day, concentrated using a Amicon Ultra centrifugal filter (molecular weight cut-off at 100 kDa, Millipore) and subjected to a Superdex 200 Increase 10/300 GL column (GE Healthcare) pre-equilibrated with buffer containing 20 mM HEPES, pH 7.5, 100 mM NaCl, 2 mM MgCl_2_, 100 μM TCEP, 10 μM GHRH and 0.001% (w/v) digitonin (Anatrace). The monomeric fractions of the SV1–G_s_ complex were collected and concentrated to 10-20 mg/mL for cryo-EM examination.

### Cryo-EM data collection and image processing

The freshly purified complexes (3.0 μL) at a final concentration of 17 mg/mL were applied to glow-discharged holey carbon grids (Quantifoil R1.2/1.3, 200 mesh), and subsequently vitrified using a Vitrobot Mark IV (ThermoFisher Scientific). Cryo-EM images were collected on a Titan Krios microscope (FEI) equipped with a K3 Summit direct electron detector (Gatan, Inc.) in the Cryo-Electron Microscopy Research Center at Shanghai Institute of Materia Medica, Chinese Academy of Sciences. A total of 7,344 movies for the *apo* SV1–G_s_ complex were automatically acquired using SerialEM (66) in super-resolution counting mode at a pixel size of 0.5225Å and with a defocus values ranging from -1.2 to -2.2 μm. Movies with 36 frames each were collected at a dose of 25 electrons per pixel per second over an exposure time of 3.2 s, resulting in an accumulated of dose of 73 electrons perÅ^2^ on sample. Image stacks of the *apo* SV1–G_s_ complex were aligned using MotionCor 2.1 (67). Contrast transfer function (CTF) parameters were estimated by Gctf v1.18 (68). The following data processing was performed using RELION-3.0-beta2 (69). Automated particle selection using Gaussian blob detection produced 4,949,167 particles from 7,344 micrographs. The particles were subjected to reference-free 2D classification to discard fuzzy particles, resulting in 2,298,424 particles for further processing. The map of GHRH-GHRHR-G_s_ complex (18) (EMD-30505) low-pass filtered to 60Å was used as the reference map for 3D classification, generating one well-defined subset with 680,397 particles. Further 3D classifications focusing the alignment on the complex produced three good subsets accounting for 377,241 particles, which were subsequently subjected to 3D refinement, CTF refinement, and Bayesian polishing. The final refinement generated a map with an indicated global resolution of 2.6Å at a Fourier shell correlation (FSC) of 0.143. For the GHRH– SV1–G_s_ complex, images were collected on a Titan Krios electron microscope (ThermoFisher Scientific) operating at 300 kV accelerating voltage, at a calibrated magnification of 130,000×, using a K2 Summit direct electron camera (Gatan) in counting mode with a Gatan Quantum energy filter. Movies were taken in EFTEM nanoprobe mode, with a 50 μm C2 aperture, corresponding to a magnified pixel size of 2.08Å on the specimen level. In total, 2397 movies were obtained with a defocus range of -1.2 to -2.2 μm. An accumulated dose of 80 electrons per Å^2^ was fractionated into a movie stack of 36 frames. Dose-fractionated image stacks were subjected to beam-induced motion correction using MotionCor2.1(67). A sum of all frames, filtered according to the exposure dose, in each image stack was used for further processing. Contrast transfer function parameters for each micrograph were determined by Gctf v1.06(68). Particle selection, 2D and 3D classifications were performed using RELION-3.0-beta2(69). Auto-picking yielded 1,632,591 particle projections that were subjected to reference-free 2D classification to discard false positive particles or particles categorized in poorly defined classes, producing 942,827 particle projections for further processing. This subset of particle projections was subjected to a round of maximum-likelihood-based 3D classifications with a pixel size of 2.08Å, resulting in one well-defined subsets with 612,594 projections. Further 3D classifications with mask on the complex produced one good subset accounting for 391,236 particles, which were subsequently subjected to a round of 3D classifications with mask on the receptor. A selected subset containing 277,500 projections was then subjected to 3D refinement and Bayesian polishing with a pixel size of 1.04Å. The final map has an indicated global resolution of 3.29Å at a FSC of 0.143.

### Model building and refinement

The final density maps of GHRH–SV1–G_s_ and *apo* SV1–G_s_ were automatically post-processed using DeepEMhancer (70) to improve the EM map quality before model building. For both structures, the initial GHRH, SV1, and G_s_ heterotrimer models were taken from the GHRH–GHRHR–G_s_–Nb35 complex (18) (PDB number: 7CZ5) and mini-G_s_ heterotrimer was taken from the GPR52–mini-G_s_ complex (71) (PDB number: 6LI3). All models were fitted into the EM density using UCSF Chimera (72), followed by iterative manual adjustment in COOT (73) according to side-chain densities. The models were then subjected to ISOLDE (74) for further rebuilding and finalized using real-space refinement in PHENIX (75). The final model statistics for both structures were validated using ‘comprehensive validation (cryo-EM)’ in PHENIX and provided in the supplementary information (*SI Appendix* Table S3). All structural figures were prepared using Chimera (72), Chimera X (76) and PyMOL (https://pymol.org/2/).

### cAMP accumulation assay

GHRH-stimulated cAMP accumulation was measured by a LANCE Ultra cAMP kit (PerkinElmer). Briefly, HEK293T cells (24 h after transfection with SV1 or GHRHR) or prostate cancer cells were digested by 0.2% (w/v) EDTA and washed once with Dulbecco’s phosphate buffered saline (DPBS). Cells were then resuspended with stimulation buffer (Hanks’ balanced salt solution (HBSS) supplemented with 5 mM HEPES, 0.5 mM IBMX and 0.1% (w/v) BSA, pH 7.4) to a density of 0.6 million cells per mL and added to 384-well white plates (3,000 cells per well). Different concentrations (5 μL) of GHRH were then added and the stimulation lasted for 30 min at RT. The reaction was stopped by adding 5 μL Eu-cAMP tracer and 5 μL ULight-anti-cAMP. After 1 h RT incubation, TR-FRET signals (excitation: 320 nm, emission: 615 nm and 665 nm) were measured by an Envision plate reader (PerkinElmer). cAMP concentrations were interpolated by a standard curve.

### G protein NanoBiT assay

HEK293T cells were seeded at a density of 30,000 cells per well into 96-well culture plates pretreated with poly-D-lysine hydrobromide. After incubation for 24 h to reach 70%-80% confluence, the cells were transiently transfected with GHRHR or SV1, Gα_s_-LgBiT, Gβ1, Gγ2-SmBiT at a 2:1:5:5 mass ratio. Twenty-four h after transfection, cells were washed once and incubated for 30 min at 37 °C with HBSS buffer (pH 7.4) supplemented with 0.1% BSA and 10 mM HEPES. They were then reacted with coelenterazine H (5 μM) for 1 h at RT. Luminescence signals were measured using a Envision plate reader at 15 s intervals (25°C). Briefly, following the baseline reading for 2.5 min, GHRH was added and the reading continued for 25 min. Data were corrected to baseline and vehicle treated samples. The area-under-the-curve (AUC) across the time-course response curve was determined and normalized to the WT GHRHR which was set to 100%.

### β-arrestin 1/2 recruitment assay

HEK293T cells were seeded at a density of 30,000 cells per well into 96-well culture plates pretreated with poly-D-lysine hydrobromide. After incubation for 24 h to reach 70%-80% confluence, the cells were transiently transfected with HA-Flag-GHRHR-Rluc8 or HA-Flag-SV1-Rluc8 and β-arrestin 1/2-Venus at a 1:9 mass ratio using Lipofectamine 3000 reagent (Invitrogen) and cultured for another 24 h. Thereafter, cells were washed once and incubated for 30 min at 37°C with HBSS buffer (pH 7.4) supplemented with 0.1% BSA and 10 mM HEPES. Five μM coelenterazine h (YEASEN Biotechnology) was then added and incubated for 5 min in the dark. The bioluminescence resonance energy transfer (BRET) signals were detected with an Envision plate reader by calculating the ratio of emission at 535 nm over emission at 470 nm. A 1.5 min baseline of BRET measurements was taken before the addition of GHRH and BRET signal was measured at 10 s intervals for further 9 min. After removing baseline and background readings by subtracting average values of the baseline measurement and average values of vehicle-treated samples, respectively, the AUC across the time-course response curve was determined and normalized to the WT GHRHR which was set to 100%.

### ERK1/2 phosphorylation assay

ERK1/2 phosphorylation was detected with the AlphaScreen SureFire ERK1/2 assay kit (PerkinElmer). Briefly, GHRHR or SV1 expressing HEK293 cells were seeded into 96-well culture plates coated with poly-D-lysine (30,000 cells/well) and grown overnight followed by deprivation of serum for at least 6 h. After incubation with FBS-free DEME medium containing 4 μM GHRHR antagonist (MIA-602) or vehicle control for 30 min at RT, ERK1/2 phosphorylation was stimulated by the addition of 1 μM GHRH (100 μL final volume) at the indicated time points. GHRH stimulation was terminated by removal of medium and addition of 30 μl of SureFire lysis buffer to each well. The plate was then agitated for 15 min. A 1:17:100 (v/v/v) dilution of AlphaScreen beads/SureFire activation buffer/SureFire reaction buffer was transferred to a white 384-well Proxiplate (8.5 μL per well) followed by addition of 5 μL lysate in diminished light. The plate was incubated in the dark at 37°C for 1 h after which the fluorescence signal was measured by an Envision plate reader, using standard AlphaScreen settings. Data were normalized to the maximal response elicited by 10% FBS for 7 min followed by normalization to the maximal response elicited by GHRH.

### Western blot

To analyze phosphorylation of ERK, HEK293T cells (24 h after transfection with SV1 or GHRHR) or prostate cancer cells were stimulated by 1 μM GHRH at the indicated time points with or without pre-treatment with 4 μM MIA-602. The cells were then lysed with RIPA lysis buffer (Sigma-Aldrich) on ice for 5 min and centrifuged at 12,000 rpm for 15 min. Protein concentration of the supernatant was determined by BCA protein assay kit (Beyotime). Proteins were loaded onto 10% sodium dodecyl sulfate-polyacrylamide gel with sodium dodecyl sulfate-loading buffer (Beyotime), separated by electrophoresis and transferred to polyvinylidene difluoride membranes (0.2 μm; Merck Millipore). The membranes were blocked with 5% BSA in TBST buffer for 2 h at RT and incubated overnight at 4°C with primary antibodies against ERK1/2 (1:1000; 9102s, Cell Signaling Technology), pERK1/2 (1:1000; 9101s, Cell Signaling Technology), β-tubulin (1:1000; 2146s, Cell Signaling Technology), GHRHR (1:800; ab76263, Abcam), flag (1:800; F3165, Sigma). After washing 3 times with TBST buffer, the membranes were incubated with secondary antibodies (1:10,000; Cell Signaling Technology) for 1 h at RT. Protein bands were visualized by ECL Plus (Bio-Rad). Densitometric analysis was then performed to determine the relative expression of target proteins normalized to β-tubulin or GAPDH. ERK1/2 phosphorylation was estimated as a ratio of pERK over total ERK.

### Cell cycle analysis

Cells were added at density of 3 × 10^5^/well to 12-well plates and treated with GHRH at indicated concentrations for 6 h. Following the treatment, the cells were digested by trypsin and collected by centrifugation. Cell pellets were washed with PBS and fixed with 70% (v/v) ethanol at 4°C overnight. Cells were again collected by centrifugation and washed twice with PBS. They were then resuspended in PBS containing: 0.2% Triton X-100, 50 μg/mL propidium Iodide (PI) and 100 mg/mL RNase A, and incubated for 30 min at 4°C in the dark. After incubation, fluorescence intensity was measured with a NovoCyte flow cytometer (Acea Biosciences). Cells were gated for PI staining and cell accumulation in G_1_, S and G_2_/M phases were calculated using the NovoExpress software.

### Molecular dynamics simulation

Previous structural studies (77-79) have reported three β-arrestin 1-bound GPCR complex structures (PDB numbers: 6TKO, 6PWC and 6UP7) but that of β-arrestin 2-bound has yet to be revealed. Thus, all the three β-arrestin 1-bound structures were used to build models. The structure of GHRH-SV1-G_s_ complex and β-arrestin 1 models were aligned to the published β-arrestin 1-bound GPCR complex structures to construct GHRH/SV1/β-arrestin-1 complex models. The missing backbone and side chains were added. Similarly, the GHRH/GHRHR/β-arrestin-1 complex models were built using the cryo-EM structure of WT GHRHR (18) (PDB code: 7CZ5). To build a simulation system, we placed the complex model into a 1-palmitoyl-2-oleoyl-sn-glycero-3-phosphocholine lipid bilayer. The lipid embedded complex model was solvated in a periodic boundary condition box (100Å × 100Å × 185Å) filed with TI3P water molecules and 0.15 M KCl using CHARMM-GUI (80, 81). Each system was replicated to perform two independent simulations. On the basis of the CHARMM36m all-atom force field (82, 83), MD simulations were conducted using GROMACS 5.1.4 (84, 85). After 100 ns equilibration, the β-arrestin models built based on β-arrestin 1 binding to β1-adrenoceptor (PDB number: 6TKO) produced stable conformations. Thus, a 1 μs production run was carried out for each simulation of these models. All productions were conducted in the NPT ensemble at temperature of 303.15 K and a pressure of 1 atm. Temperature and pressure were controlled using the velocity-rescale thermostat (86) and the Parrinello-Rahman barostat with isotropic coupling (87), respectively. Equations of motion were integrated with a 2 fs time step, the LINCS algorithm was used to constrain bond length (88). Nonbonded pair lists were generated every 10 steps using distance cut-off of 1.4 nm. A cut-off of 1.2 nm was used for Lennard-Jones (excluding scales 1-4) interactions, which were smoothly switched off between 1 and 1.2 nm. Electrostatic interactions were computed using particle-mesh-Ewald algorithm with a real-space cut-off of 1.2 nm (89). The last 200 ns trajectory of each simulation was used to calculate average values. The distance between two residues is the minimal distance between non-hydrogen atoms from two different residues; the distance between the non-hydrogen atoms from the same residue was excluded. The backbone distance between two motifs is the minimal distance between backbone atoms from two different defined motifs; the distance between the backbone atoms from the same motif was excluded. The frequency of a particular residue interacting with a peptide was calculated by counting how many times this residue interacts with the peptide in the simulation snapshots. The interaction is defined by the non-hydrogen atom distance between the residue and the peptide using 4Å as cut-off. The interacting frequency value indicates the stability of a particular residue-peptide interaction. A large interacting frequency indicates a stable interaction.

### Statistical analysis

All functional study data were analyzed using Prism 7 (GraphPad) and presented as means ± S.E.M. from at least three independent experiments. Concentration-response curves were evaluated with a three-parameter logistic equation. The significance was determined with either two-tailed Student’s *t*-test or one-way ANOVA, and P < 0.05 is considered statistically significant.

## Acknowledgments

We are indebted to Zhijie Liu and Hua Tian for valuable discussions. This work was partially supported by National Natural Science Foundation of China 81872915 (M.-W.W.), 82073904 (M.-W.W.), 81773792 (D.Y.), 81973373 (D.Y.) and 21704064 (Q.Z.); National Science & Technology Major Project of China–Key New Drug Creation and Manufacturing Program 2018ZX09735–001 (M.-W.W.) and 2018ZX09711002–002–005 (D.Y.); the National Key Basic Research Program of China 2018YFA0507000 (M.-W.W.); Shanghai Municipal Science and Technology Major Project 2019SHZDZX02 (H.E.X.); Ministry of Science and Technology of China Major Project XDB08020303 (H.E.X.); Novo Nordisk-CAS Research Fund grant NNCAS-2017–1-CC (D.Y.); The Youth Innovation Promotion Association of CAS 2018319 (X.C); The Young Innovator Association of CAS Enrollment (H.M. and L.H.Z.) and SA-SIBS Scholarship Program (L.H.Z. and D.Y.). The cryo-EM data were collected at Cryo-Electron Microscopy Research Center, Shanghai Institute of Materia Medica.

## Notes

### Competing Interest Statement

The authors have declared no competing interest.

