## Supporting Information for "Constitutive signal bias mediated by the human GHRHR splice variant 1"

[1]Department of Pharmacology, School of Basic Medical Sciences, Fudan University, Shanghai 200032, China; [2]School of Pharmacy, Fudan University, Shanghai 201203, China; [3]The CAS Key Laboratory of Receptor Research, Shanghai Institute of Materia Medica, Chinese Academy of Sciences, Shanghai 201203, China; [4]School of Life Science and Technology, ShanghaiTech University, Shanghai 201210, China; [5]University of Chinese Academy of Sciences, Beijing 100049, China; [6]School of Artificial Intelligence and Automation, Huazhong University of Science and Technology, Wuhan 430074, China; [7]Department of Biophysics and Department of Pathology of Sir Run Run Shaw Hospital, Zhejiang University School of Medicine, Hangzhou 310058, China; [8]The National Center for Drug Screening, Shanghai Institute of Materia Medica, Chinese Academy of Sciences, Shanghai 201203, China; [9]Eye and ENT Hospital, Fudan University, Shanghai 200031, China; [10]State Key Laboratory of Drug Research and Drug Discovery and Design Center, Shanghai Institute of Materia Medica, Chinese Academy of Sciences, 201203, Shanghai, China; [11]These authors contributed equally to this work.

***Correspondence:** H. Eric Xu, Xi Cheng, Dehua Yang, Ming-Wei Wang

**This PDF file includes:**

Figures S1 to S9
Tables S1 to S4

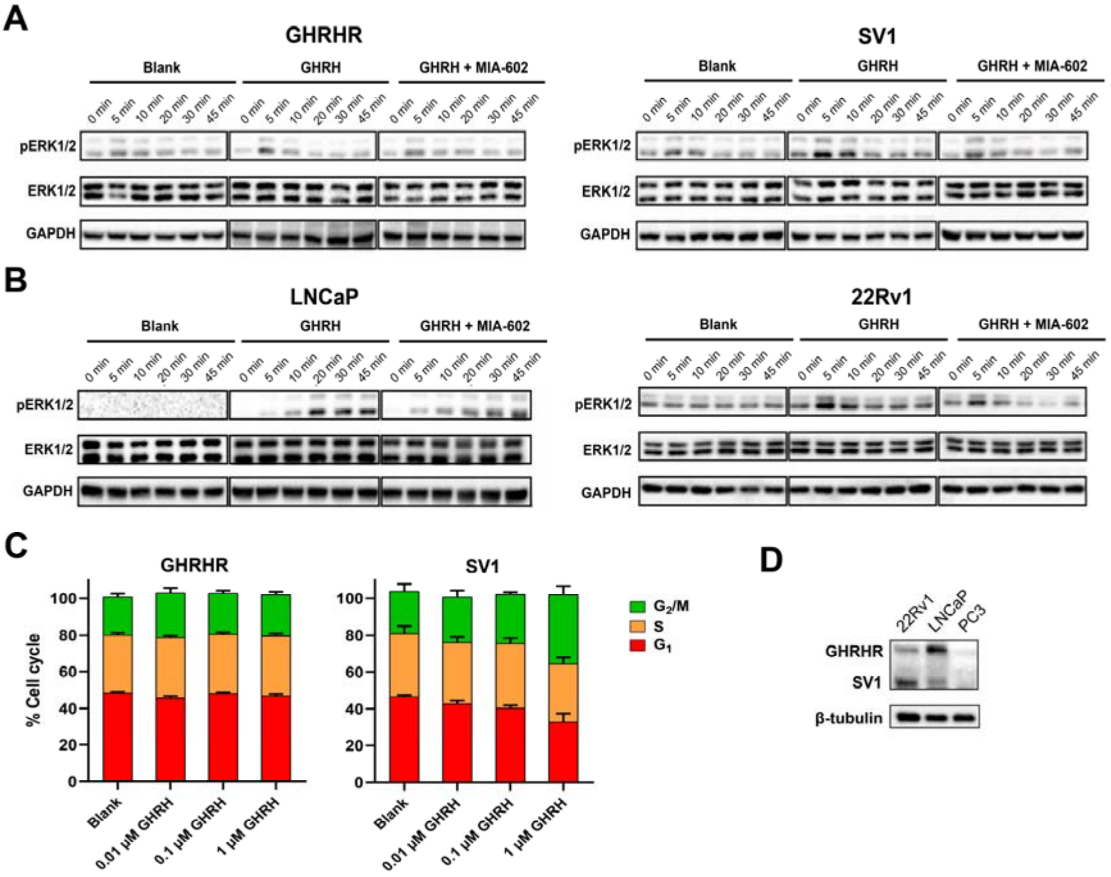


**Figure S1. GHRH-induced pERK1/2 signaling mediated by SV1 and GHRHR and concurrent cell cycle change.** (*A*, *B*) Representative time-course signaling of ERK1/2 monitored by immunoblotting of the total ERK1/2 and phosphorylated ERK1/2 (pERK1/2). The assay was initiated by 1 μM GHRH and inhibition was achieved by 4 μM MIA-602 in HEK293T cells expressing GHRHR or SV1 (*A*) and prostate cancer cell lines (*B*). (*C*) Cell distribution in $G_1$, S and $G_2$/M phases after treatment of different concentrations of GHRH. Data shown are means ± S.E.M. of at least three independent experiments (*n* = 3-5) performed in duplicate. (*D*) Expression of GHRHR and SV1 in prostate cancer cell lines.

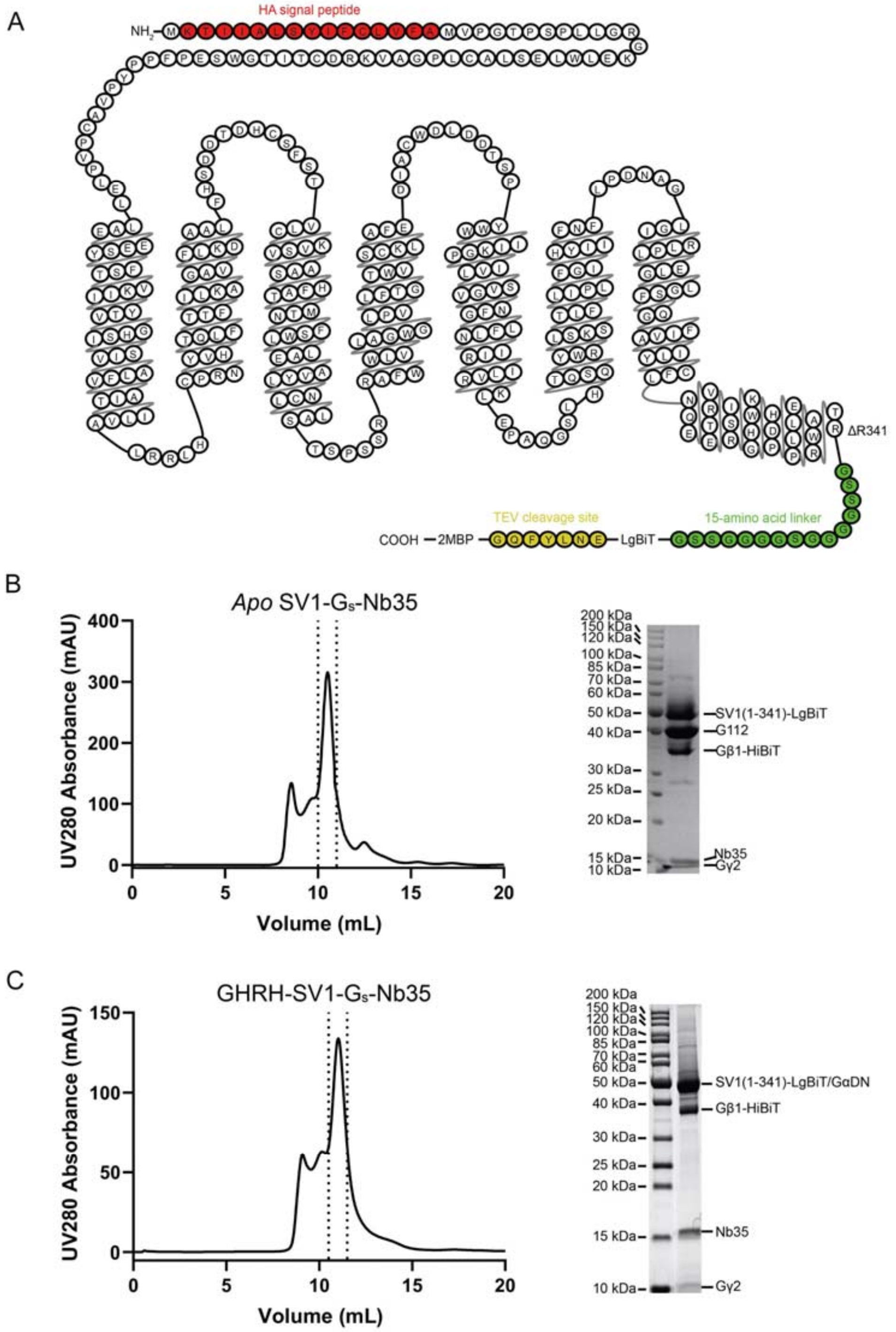


**Figure S2. Purification and characterization of the SV1–G$_s$–Nb35 complex.** (*A*) Schematic of the HA-SV1(1-341)-15AA-LgBiT-TEV-2MBP construct used in cryo-EM study. The HA signal peptide (red), 15-amino acid (AA) linker (green), Tev cleavage site (yellow) and R341 truncation site are highlighted and indicated. (*B*, *C*) Size-exclusion chromatography elution profile and corresponding SDS-PAGE gel of the *apo* SV1–G$_s$–Nb35 (*B*) and GHRH–SV1–G$_s$–Nb35 (*C*) complexes. G112 is an engineered G$\alpha_s$ protein.

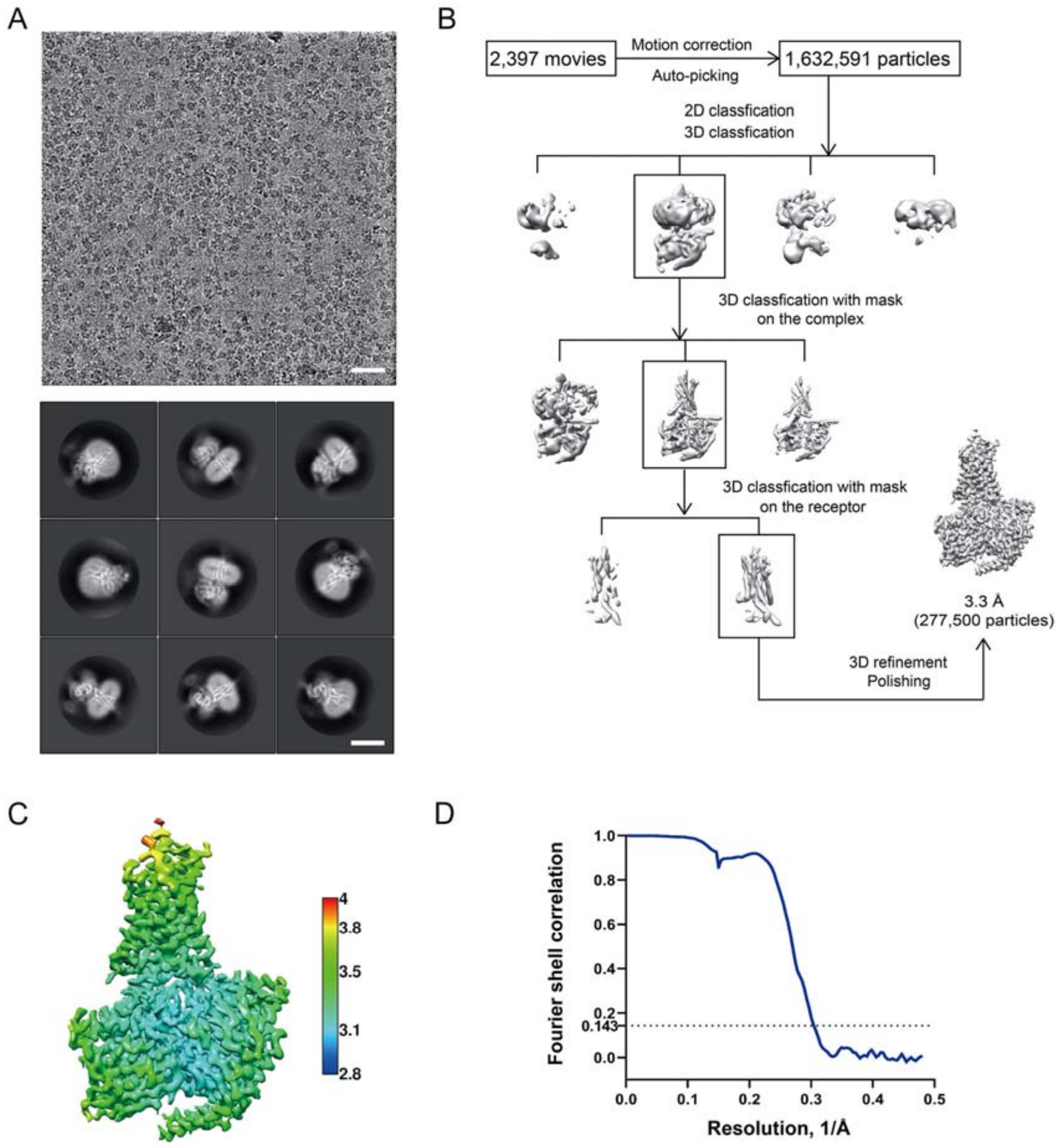


**Figure S3. Cryo-EM data processing and validation of the GHRH–SV1–$G_s$–Nb35 complex.** (*A*) Representative cryo-EM micrograph (scale bar: 40 nm) and two-dimensional class averages (scale bar: 5 nm). (*B*) Flow chart of cryo-EM data processing. (*C*) Local resolution distribution map of the complex. (*D*) Fourier shell correlation (FSC) curve of the overall refined receptor.

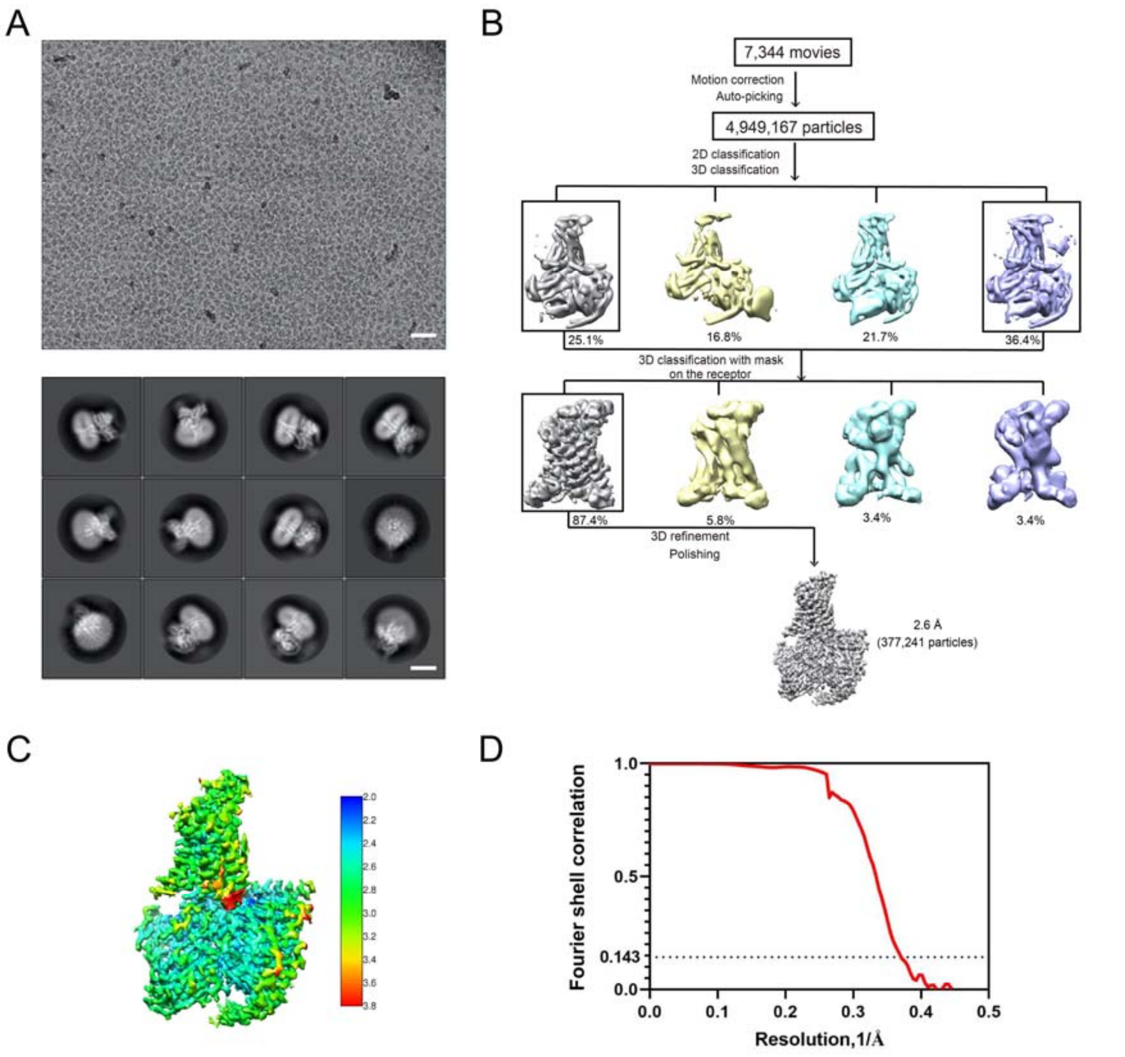


**Figure S4. Cryo-EM data processing and validation of the *apo* SV1–$G_s$–Nb35 complex.** (*A*) Representative cryo-EM micrograph (scale bar: 40 nm) and two-dimensional class averages (scale bar: 5 nm). (*B*) Flow chart of cryo-EM data processing. (*C*) Local resolution distribution map of the complex. (*D*) FSC curves of the overall refined receptor.

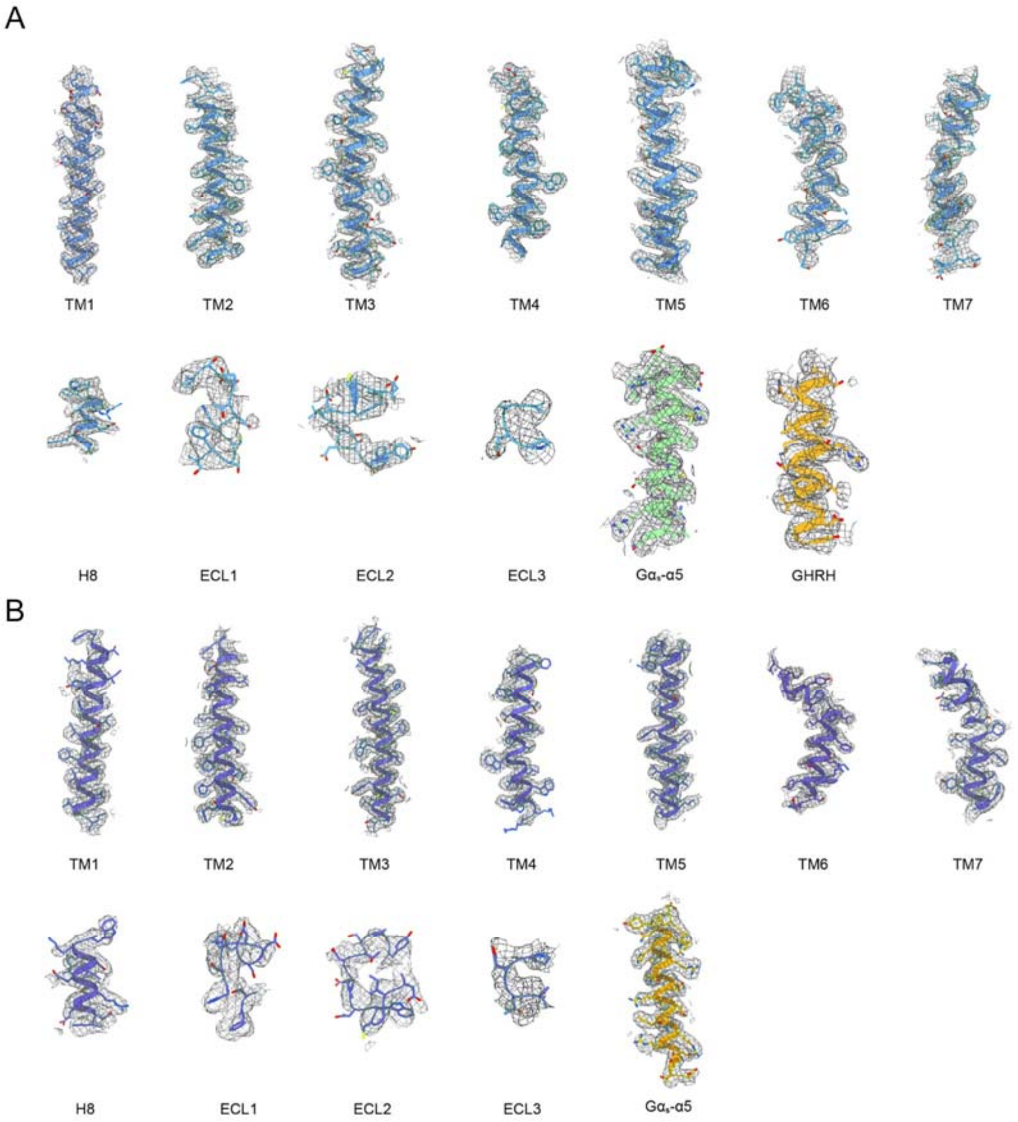


**Figure S5. Cryo-EM density map of the GHRH–SV1–$G_s$ and the *apo* SV1–$G_s$ structures.** (*A*) Cryo-EM density map and model of the GHRH–SV1–$G_s$ structure are shown for all seven-transmembrane (TM) α-helices, ECLs 1-3, helix 8 (H8) of SV1, GHRH, Gα and helix α5. (*B*) Cryo-EM density map and model of the *apo* SV1–$G_s$ structure are shown for all 7-TM α-helices, ECLs 1-3, helix 8 of SV1, Gα and helix α5.

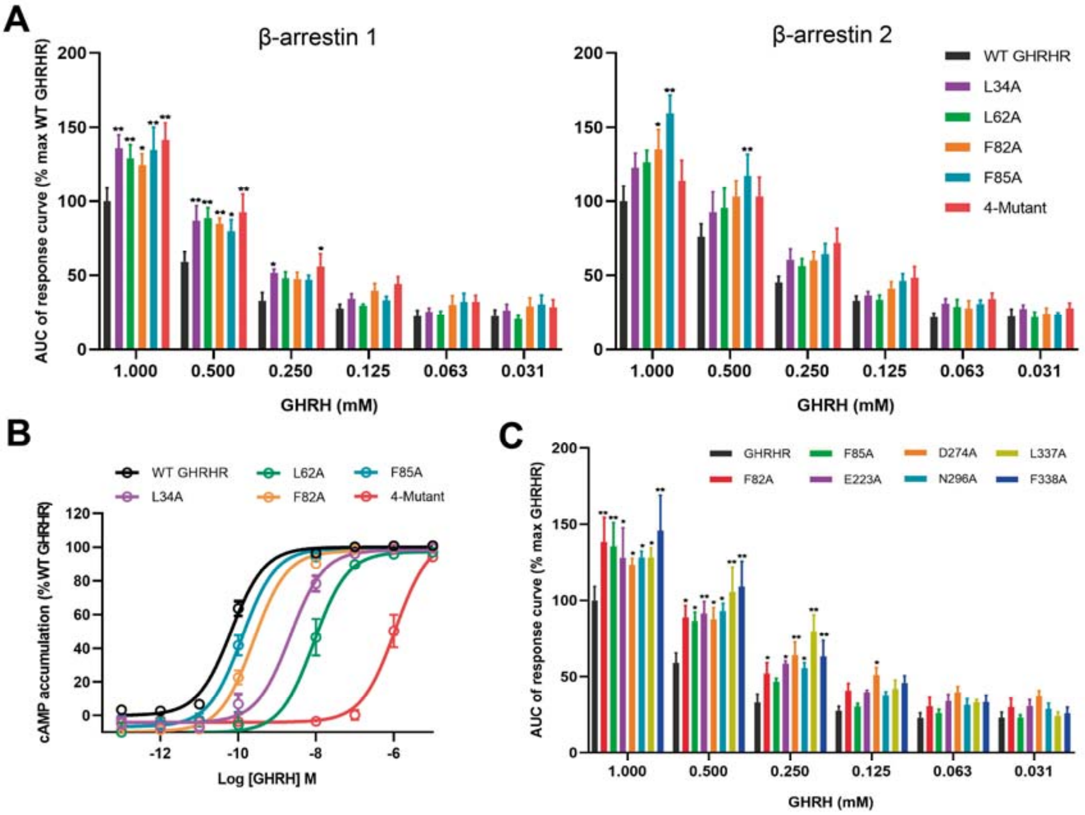


**Figure S6. Residues of GHRHR responsible for biased signaling.** (*A*) Mutations in the extracellular domain (ECD) reduced the cAMP response of GHRHR. (*B*) Mutations in the ECD affect β-arrestin 1/2 recruitment by GHRHR. 4-Mutant, single-point GHRHR mutation made simultaneously at 4 residues, L34A, L62A, F82A and F85A. (*C*) β-arrestin 1 recruitment by GHRHR and its mutants. Data shown are means ± S.E.M. of five independent experiments ($n$ = 5) performed in quadruplicate or duplicate, respectively; *$P$ < 0.05, **$P$ < 0.01. WT, wild-type; max, maximum.

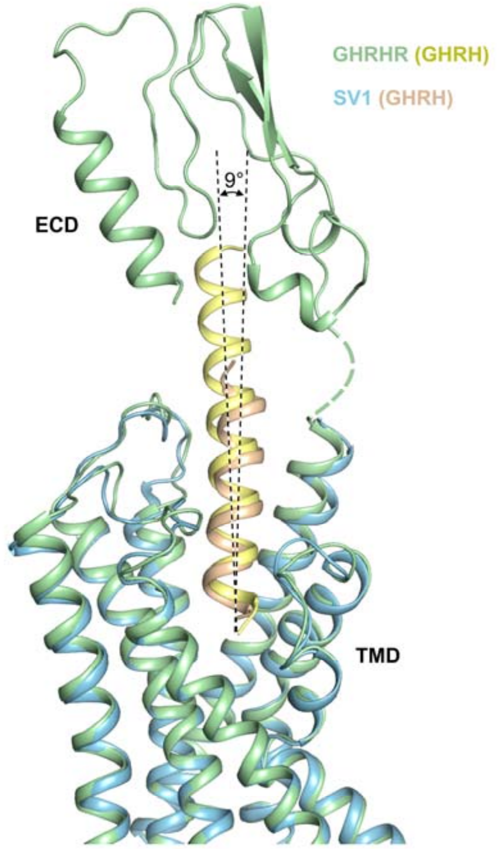


**Figure S7. Comparison between the cryo-EM structures of GHRH–SV1–$G_s$ and GHRH–GHRHR–$G_s$ complexes at the extracellular side.** Receptors and GHRH are shown in cartoon: GHRHR is colored in green, SV1 in blue, GHRH in wheat and yellow. $G_s$ is omitted for clarity. ECD, extracellular domain; TMD, transmembrane domain.

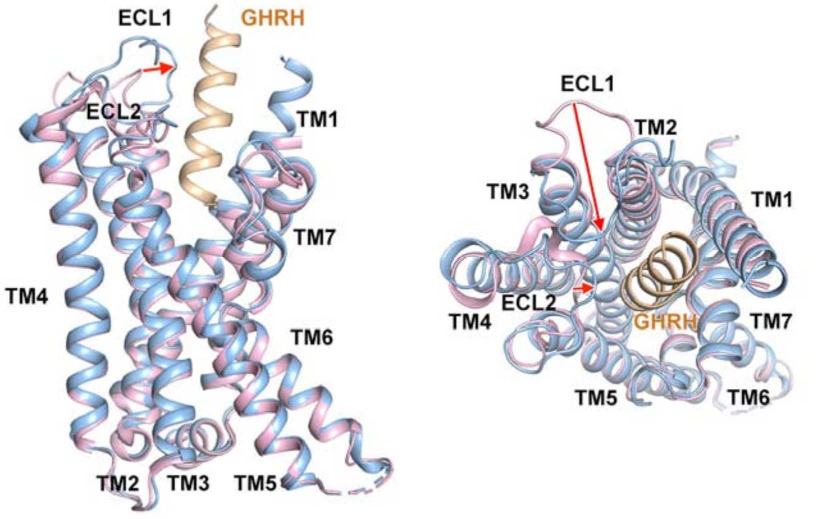


**Figure S8. Comparison of the GHRH–SV1–$G_s$ complex structure with that of the *apo* SV1–$G_s$ complex.** Both side (left) and top (right) views are displayed. Receptors and GHRH are shown in cartoon. In the GHRH–SV1–$G_s$ complex structure, SV1 is colored in blue and GHRH is in wheat. In the *apo* SV1–$G_s$ complex structure, SV1 is colored in pink. $G_s$ is omitted for clarity.

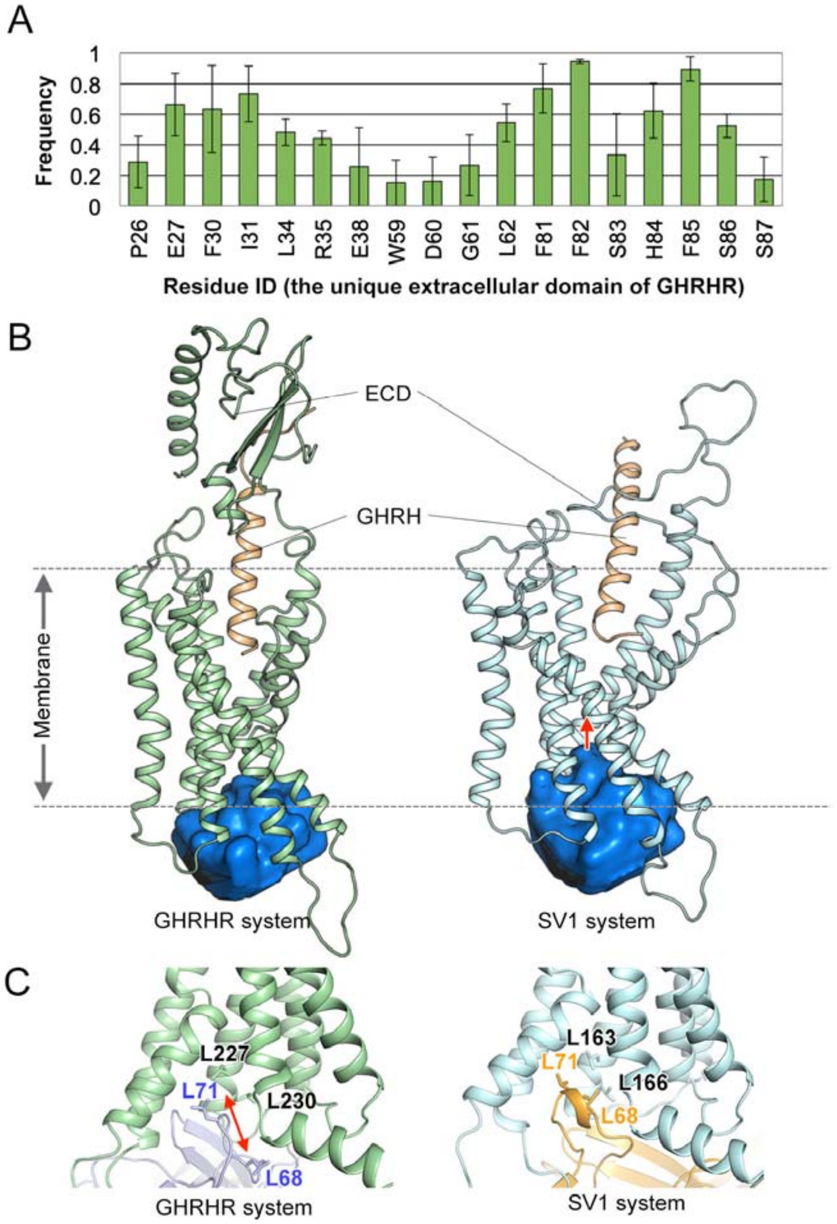


**Figure S9. Extracellular and intracellular interactions of GHRHR and SV1.** (*A*) Interacting frequency between an ECD residue of GHRHR and GHRH. The interacting frequency value indicates the stability of a particular residue-peptide interaction. A large interacting frequency implies a stable interaction. (*B*) Representative simulation snapshots from GHRHR system (left) and SV1 system (right). Receptors and GHRH are shown in cartoon: GHRHR is colored in green, SV1 in blue, GHRH in wheat. β-arrestin 1 is omitted for clarity. Arrestin-binding pockets are shown in surface depict. (*C*) A representative simulation snapshot showing key interactions of GHRHR (green) and SV1 (blue) at intracellular side. Key residues are shown as sticks.

**Table S1**. Effects of SV1 on GHRH-induced $G_s$ activation.

| **Plasmid** | **$pEC_{50}$** | **$E_{max}$ (% WT GHRHR)** |
|---|---|---|
| GHRHR/$G_s$ | 8.17 ± 0.17 | 84.99 ± 3.32 |
| SV1/$G_s$ | 5.63 ± 0.36* | 56.67 ± 13.79* |
| Vector/$G_s$ | NA | NA |

G protein NanoBiT data were analyzed using a three-parameter logistic equation to determine $pEC_{50}$ and $E_{max}$ values. $pEC_{50}$ is the negative logarithm of the molar concentration of agonist that induced half the maximal response. $E_{max}$ is expressed as a percentage of GHRHR/$G_s$ response. All values are means ± S.E.M. of five independent experiments ($n$ = 5) conducted in duplicate. One-way ANOVA was used to determine statistical significance (*$P < 0.05$). WT, wild-type; NA, not active.

**Table S2**. Effects of residue mutation or truncation in the ECD of GHRHR on GHRH-induced cAMP accumulation.

| Mutant | $pEC_{50}$ | $E_{max}$ (% WT GHRHR) | Cell surface expression (% WT GHRHR) | ΔLog ($\tau/K_A$) |
|---|---|---|---|---|
| WT GHRHR | 10.38 ± 0.07 | 100.5 ± 1.81 | 100 | 0 ± 0.05 |
| SV1 | 6.43 ± 0.06*** | 104.85 ± 2.20 | 31.37 ± 6.83 | -2.98 ± 0.04*** |
| GHRHR Δ89 | 6.27 ± 0.06*** | 106.29 ± 2.48 | 49.46 ± 8.69 | -3.28 ± 0.03*** |
| GHRHR ΔECD | 5.53 ± 0.07*** | 111.27 ± 3.65 | 209.26 ± 5.61 | -4.55 ± 0.04*** |
| GHRHR Δ32 | 6.55 ± 0.05*** | 105.06 ± 1.93 | 79.27 ± 3.78 | -3.23 ± 0.04*** |
| GHRHR Δ42 | 6.95 ± 0.04*** | 104.23 ± 1.53 | 91.80 ± 4.81 | -2.86 ± 0.02*** |
| GHRHR Δ52 | 6.62 ± 0.04*** | 105.63 ± 1.62 | 40.54 ± 5.0 | -2.83 ± 0.03*** |
| GHRHR Δ62 | 6.34 ± 0.05*** | 103.91 ± 2.19 | 31.30 ± 6.55 | -3.05 ± 0.03*** |
| GHRHR Δ72 | 6.44 ± 0.06*** | 104.35 ± 2.57 | 35.73 ± 7.58 | -2.98 ± 0.04*** |
| GHRHR Δ82 | 6.23 ± 0.06*** | 104.32 ± 2.51 | 31.60 ± 5.79 | -3.14 ± 0.03*** |
| GHRHR Δ92 | 6.25 ± 0.06*** | 105.98 ± 2.38 | 45.34 ± 5.32 | -3.78 ± 0.03*** |
| GHRHR Δ102 | 6.09 ± 0.05*** | 107.29 ± 2.15 | 121.17 ± 12.35 | -3.55 ± 0.02*** |
| GHRHR Δ112 | 5.93 ± 0.05*** | 106.27 ± 2.34 | 54.92 ± 7.98 | -3.65 ± 0.02*** |
| L34A | 8.67 ± 0.11*** | 100.7 ± 2.23 | 94.27 ± 12.02 | -0.78 ± 0.06*** |
| L62A | 8.06 ± 0.10*** | 97.99 ± 2.00 | 62.40 ± 5.34 | -1.03 ± 0.04*** |
| F82A | 9.61 ± 0.49*** | 99.77 ± 1.36 | 102.74 ± 12.13 | -0.53 ± 0.05*** |
| F85A | 9.91 ± 0.06** | 100.24 ± 1.69 | 96.75 ± 10.38 | -0.27 ± 0.04*** |
| L34A-L62A-F82A-F85A | 5.97 ± 0.09*** | 105.23 ± 5.52 | 77.50 ± 6.89 | -4.07 ± 0.04*** |

cAMP accumulation data were analyzed using a three-parameter logistic equation to determine $pEC_{50}$ and $E_{max}$ values. $pEC_{50}$ is the negative logarithm of the molar concentration of agonist that induced half the maximal response. $E_{max}$ for mutants is expressed as a percentage of the WT GHRHR. Data were analyzed by nonlinear regression using the operational model equation to determine the logR values ($\log\tau/K_A$, *i.e.,* logarithm of the transduction ratio). $\tau$ is the efficacy value of the agonist and was corrected by cell surface expression of the receptor. $K_A$ is dissociation constant. Changes in transduction ratio (ΔlogR) were calculated to determine the relative effectiveness of the mutants. All values are means ± S.E.M. of at least three independent experiments ($n$ = 3-5) conducted in quadruplicate. Statistical analysis was carried out by comparing the control responses in the WT GHRHR. **, $P < 0.01$ and ***, $P < 0.001$, determined by one-way ANOVA.

**Table S3.** Cryo-EM data collection, model refinement and validation statistics.

| **Data collection and processing** | **GHRH-SV1-$G_s$-Nb35** | **SV1-$G_s$-Nb35** |
|---|---|---|
| Magnification | 130,000 | 130,000 |
| Voltage (kV) | 300 | 300 |
| Electron exposure (e–/Å$^2$) | 80 | 73 |
| Defocus range (μm) | -1.2 to -2.2 | -1.5 to -2.5 |
| Pixel size (Å) | 1.04 | 1.045 |
| Symmetry imposed | C1 | C1 |
| Initial particle images (no.) | 1,632,591 | 4,949,167 |
| Final particle images (no.) | 277,500 | 377,241 |
| Map resolution (Å) | 3.29 | 2.60 |
| FSC threshold | 0.143 | 0.143 |
| Map resolution range (Å) | 2.9-4.6 | 2.2-4.0 |
| **Refinement** | | |
| Initial model used (PDB code) | 7CZ5 | 7CZ5 |
| Model resolution (Å) | 3.4 | 3.1 |
| FSC threshold | 0.5 | 0.5 |
| Map sharpening B factor (Å$^2$) | -104.32 | -82.56 |
| Model composition | | |
| Non-hydrogen atoms | 8139 | 8047 |
| Protein residues | 1029 | 1019 |
| B factors (Å$^2$) | | |
| Protein | 57.85 | 93.64 |
| Root mean square deviation | | |
| Bond lengths (Å) | 0.004 | 0.004 |
| Bond angles (°) | 0.643 | 0.679 |
| Validation | | |
| MolProbity score | 1.52 | 1.39 |
| Clash score | 4.39 | 3.50 |
| Poor rotamers (%) | 0.23 | 0.0 |
| Ramachandran plot | | |
| Favored (%) | 95.63 | 96.31 |
| Allowed (%) | 4.37 | 3.69 |
| Disallowed (%) | 0.0 | 0.0 |

**Table S4**. Effects of residue mutation in the ligand-binding pocket of SV1 on GHRH-induced cAMP accumulation.

| Mutant | $pEC_{50}$ | $E_{max}$ (% WT GHRHR) | Cell surface expression (% WT GHRHR) | ΔLog ($\tau/K_A$) |
|---|---|---|---|---|
| GHRHR | 10.83 ± 0.04 | 100 | 100 | 0.00 ± 0.09 |
| SV1 | 6.38 ± 0.06 | 101.74 ± 2.28 | 47.56 ± 3.76 | -3.93 ± 0.10 |
| F62A | 5.28 ± 0.08** | 105.64 ± 4.54 | 58.47 ± 6.68 | -4.41 ± 0.14*** |
| V65A | 5.33 ± 0.09* | 106.08 ± 4.91 | 49.35 ±1.32 | -4.08 ± 0.12 |
| K66A | 5.6 ± 0.08 | 106.82 ± 4.09 | 13.88 ± 2.07 | -4.02 ± 0.11 |
| Y69A | 5.44 ± 0.09* | 104.75 ± 4.29 | 36.69 ± 1.41 | -3.81 ± 0.10 |
| K118A | 4.61 ± 0.12*** | 92.4 ± 7.25 | 57.63 ± 4.09 | -4.51 ± 0.13*** |
| S145A | 6.34 ± 0.09 | 101.87 ± 3.5 | 37.13 ± 2.32 | -4.22 ± 0.14** |
| H146A | 5.07 ± 0.08*** | 101.06 ± 4.83 | 57.74 ± 7.57 | -4.64 ± 0.11*** |
| I225A | 4.98 ± 0.12*** | 76.03 ± 5.48*** | 23.03 ± 2.05 | -3.85 ± 0.13 |
| L290A | 5.58 ± 0.08 | 109.6 ± 3.9 | 34.40 ± 7.20 | -4.56 ± 0.09*** |
| L294A | 5.32 ± 0.1* | 106.26 ± 5.25 | 35.47 ± 1.85 | -4.96 ± 0.12*** |

cAMP accumulation data were analyzed using a three-parameter logistic equation to determine $pEC_{50}$ and $E_{max}$ values. $pEC_{50}$ is the negative logarithm of the molar concentration of agonist that induced half the maximal response. $E_{max}$ for mutants is expressed as a percentage of the WT GHRHR. Data were analyzed by nonlinear regression using the operational model equation to determine the logR values ($\log\tau/K_A$, *i.e.*, logarithm of the transduction ratio). $\tau$ is the efficacy value of the agonist and was corrected by cell surface expression of the receptor. $K_A$ is dissociation constant. Changes in transduction ratio (ΔlogR) were calculated to determine the relative effectiveness of the mutants. All values are means ± S.E.M. of four independent experiments ($n$ = 4) conducted in quadruplicate. Statistical analysis was carried out by comparing the control responses in the WT-SV1. *, $P < 0.05$, **, $P < 0.01$ and ***, $P < 0.001$, determined by one-way ANOVA.
